# Stabilization of a Protein by a Single Halogen-Based Aromatic Amplifier

**DOI:** 10.1101/2022.07.25.501420

**Authors:** Krystel El Hage, Nelson B. Phillips, Yen-Shan Chen, Balamurugan Dhayalan, Jonathan Whittaker, Kelley Carr, Linda Whittaker, Manijeh H. Phillips, Faramarz Ismail-Beigi, Markus Meuwly, Michael A. Weiss

**Affiliations:** Department of Chemistry, University of Basel, Klingelbergstrasse 80 CH-4056 Basel, Switzerland; Department of Biochemistry, Case Western Reserve University, Cleveland, Ohio 44106, USA; Department of Medicine, Case Western Reserve University, Cleveland, Ohio 44106, USA; Department of Biochemistry & Molecular Biology, Indiana University School of Medicine, Indianapolis, IN 46202, USA

**Author notes:** SABNP, Univ. Evry, INSERM U1204, Université Paris-Saclay, 91025 Evry, France. These authors contributed equally (Krystel El Hage, Nelson B. Phillips).

## Abstract

The utility of halogenation in protein design is investigated by a combination of quantitative atomistic simulations and experiment. The approach is applied to insulin, a small, therapeutically relevant domain amenable to simulation and semi-synthesis. In a singly halogenated aromatic ring, the simulations predicted regiospecific inductive effects to modulate multiple surrounding electrostatic (weakly polar) interactions, thereby amplifying changes in thermodynamic stability. In accordance with the simulations, stabilization of insulin is demonstrated by single halogen atoms at the ortho position of an invariant phenylalanine (2-F-PheB24, 2-Cl-PheB24 and 2-Br-PheB24; ΔΔ*G*_*u*_ = -0.5 to -1.0 kcal/mol) located at the edge of a protein crevice. Corresponding meta and para substitutions have negligible effects. The ortho-modified insulin analogs exhibit enhanced resistance to fibrillation above room temperature and retain biological activity in mammalian cells and in a rat model of diabetes mellitus. Consequently, halogen-based stabilization of insulin and other therapeutic proteins may provide a biophysical strategy to circumvent the requirement for a distribution “cold chain” in the developing world and enhance the shelf life of pharmaceutical formulations.

## Introduction

The engineering of functionally intact ultra-stable proteins, including therapeutic agents and vaccines, bears broad societal benefits in regions of the developing world where electricity and refrigeration are not consistently available. An example of a therapeutic protein susceptible to thermal degradation is provided by insulin.^1^ Here, quantitative atomistic simulations^2^ are combined with experiments for insulin analogs to demonstrate that protein stability can be markedly enhanced by a single halogen atom while maintaining biological activity. The utility of halogenic substitutions is well established in medicinal chemistry.^3^ Fluorinated functional groups are critical, for example, to the efficacy of widely prescribed small molecules such as *atorvastatin* (Lipitor^®^), an inhibitor of cholesterol biosynthesis,^4^ and *fluoxetine hydrochloride* (Prozac^®^), a selective serotonin re-uptake inhibitor.^5, 6^ Although the atomic radius of fluorine is similar to that of hydrogen (larger by only 0.27 Å), its substantial inductive effect modifies the stereo-electronic properties of these drugs, and enhances their bioactivities.^3^ Such observations have motivated the study of fluorinated amino acids in proteins^7^ and the development of “extended genetic-code” technology for their biosynthetic incorporation.^8^

Attention has largely focused on the use of multiply fluorinated aliphatic side chains (such as trifluoro-γ-CF_3_-Val, trifluoro-δ-CF_3_-Val, trifluoro-δ-CF_3_-Ile, hexafluoro-γ_1,2_-CF_3_-Val, and hexafluoro-δ_1,2_-CF_3_-Leu) to augment hydrophobicity within the cores of globular proteins. An example is provided by the stabilization of a model α-helical motif, the homodimeric coiled coil.^9^ Substitution of a fluorous interface enhances stability by 0.3 to 2.0 kcal/mol.^10-13^ This approach requires multiple fluorous substitutions, since the stability change *per fluorine atom* is small (ΔΔ*G*_u_ < 0.05 kcal/mol). Substitution of a specific Phe by *penta*-fluoro-Phe in villin headpiece (a small globular protein) within a cluster of internal aromatic rings was found to preserve its structure^14^ with a gain in stability of 0.6 kcal/mol (ΔΔ*G*_u_ = -0.12 kcal/mol per halogen atom).^15^ Such stabilization depended exquisitely on structural details as single Phe→*penta*-fluoro-Phe at six other sites in the domain were destabilizing. Although larger halogens have also been employed in medicinal chemistry (such as a chloro-aromatic substitution in chloroquine [(*RS*)-*N’*-(7-chloroquinolin-4-yl)-*N,N*-diethyl-pentane-1,4-diamine] and related anti-malarial compounds^16^) and a bromo-aromatic substitution in bromocriptine ((5′α)-2-Bromo-12′-hydroxy-5′-(2-methylpropyl)-3′,6′,18-trioxo-2′-(propan-2-yl)ergotaman^17^)), analogous applications in protein engineering have been limited.

The present study was motivated by the hypothesis that an aromatic system in a protein can act as an “amplifier” of per-halogen stabilization free energy. The rationale was that, unlike aliphatic substitutions (such as perfluoro-Leu), regiospecific inductive perturbations in a π-system may be transmitted via surrounding “weakly polar” interactions^18, 19^ to augment the resulting net gain or loss of stability. A particularly relevant system is provided by an invariant residue in insulin (Phe^B24^) whose aromatic side chain packs at the edge of the hydrophobic core^20^ at a classical receptor-binding surface (Figure 1a).^20, 21^ The biological importance of Phe^B24^ is highlighted by a clinical mutation (Ser^B24^) causing human diabetes mellitus.^22^ Lying within the C-terminal Δ-stand of the B-chain (residues B24-B28), Phe^B24^ adjoins the central α-helix (residues B9-B19). One face of the aromatic ring is exposed to solvent whereas the other sits within a shallow pocket whose floor is defined by Leu^B15^ and Cys^B19^ (Figure 1b). The rim of the pocket contains main-chain carbonyl- and amide groups at one edge and an adjoining aromatic ring (of Tyr^B26^) at another, together creating a complex electrostatic environment. Recent cryo-EM-based and crystallographic studies of the insulin receptor (IR) ectodomain^23, 24^ and its fragments^21^ have demonstrated that Phe^B24^ participates in a conformational switch in the insulin B-chain.^25^ On receptor engagement, the aromatic side chain inserts within a nonpolar binding pocket at the hormone-IR interface.^21^

**Figure 1.**
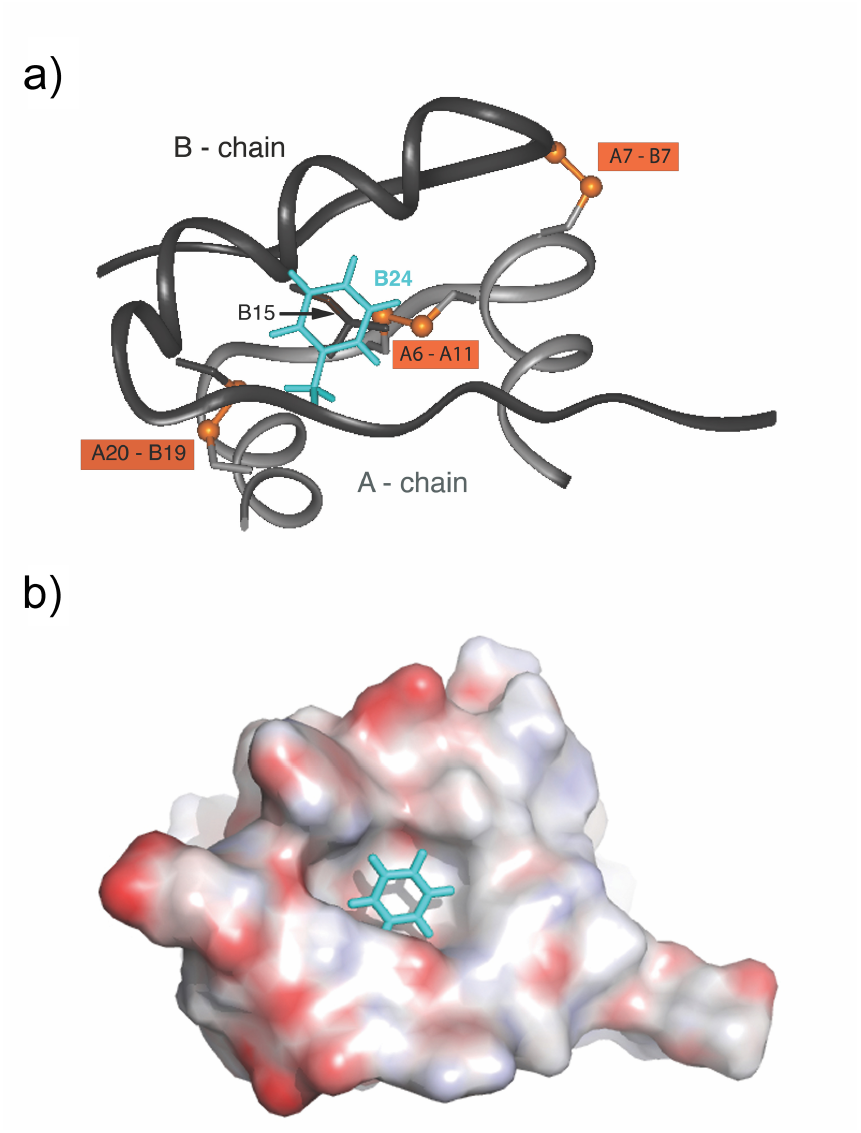
Position of Phe^B24^ within structure of an insulin monomer. **a**. Ribbon model showing position of Phe^B24^ (*aquamarine*) with respect to disulfide bridges (*tawny*). The A- and B-chains are shown in light and dark *gray*, respectively. **b**. Positioning of Phe^B24^ aromatic ring in a crevice at edge of hydrophobic core. Space-filling molecule of the remainder of the insulin molecule is color-coded by electrostatic potential (*red*, negative; and *blue*, positive).

The present study has three parts. In an effort to exploit weakly polar interactions in an asymmetric structural environment, first atomistic simulations were employed to investigate the potential role of Phe^B24^ as an “aromatic amplifier,” *i*.*e*., a mechanism to enhance the thermodynamic effects of a single halogen atom within a protein. Next, an extensive series of insulin analogs was prepared to test these predictions *<u>in vitro</u>*. Finally, biological studies were undertaken in cell culture and in a rat model of diabetes mellitus.^26^ Together, the present results provide theoretical and experimental evidence that a single halo-aromatic substitution can be as stabilizing in a protein crevice as the global replacement of aliphatic side chains by their trifluoromethyl analogs. Because an aromatic amplifier can enhance the stability while preserving biological activity, our halogen-based strategy may facilitate the global distribution and use of therapeutic proteins.

## Results and Discussion

### Halo-aromatic Substitutions Introduce Novel Weakly Polar Interactions

To evaluate halo-aryl substitutions at B24, first analyzed the electrostatic properties of the modified rings as visualized via color-coded electrostatic-surface potentials (ESP). The electrostatic potential surfaces (ESPs) of related model compounds (benzene, fluorobenzene, chlorobenzene and bromobenzene) were obtained from *ab initio* calculations and mapped at the 10^−3^ ea^-3^ isodensity surface; see Figure 2. Three physical aspects are pertinent: (i) Because halogens are electron-withdrawing substituents, such single modification lead to reorganization of electron density, creating a permanent electrostatic dipole moment directed at the halogen atom. However, relative changes in quadrupolar or higher-order moments are negligible. (ii) A halo-aromatic substitution also decreases the overall negative electrostatic potential of *π*-orbitals (as signified by the shift in color in Figure 2 from red to yellow above the ring): the larger the halogen (in the series F < Cl < Br), the weaker is the aromatic electron density (left to right in Figure 2). This would in principle decrease the strength of potential *π*−*π* interactions in protein interiors.^19^ (iii) Whereas the above two mechanisms are non-local, the larger halogen atoms provide a local electrostatic potential with a “janus” character: an electrostatically positive region (*δ*^+^) on the X-atom and along the C-X bond (X being the halogen) and an electrostatically negative region (*δ*^−^) on its equatorial flanks. The former (“*σ*-hole”) can engage in halogen bonding along the C-X bond^27^ whereas the latter can interact with sites of positive partial charge such as peptide bonds or the carboxamide functions of Asn or Gln. Such favorable electrostatic interactions are “weakly polar”^19^ akin to hydrogen bonds.

**Figure 2.**
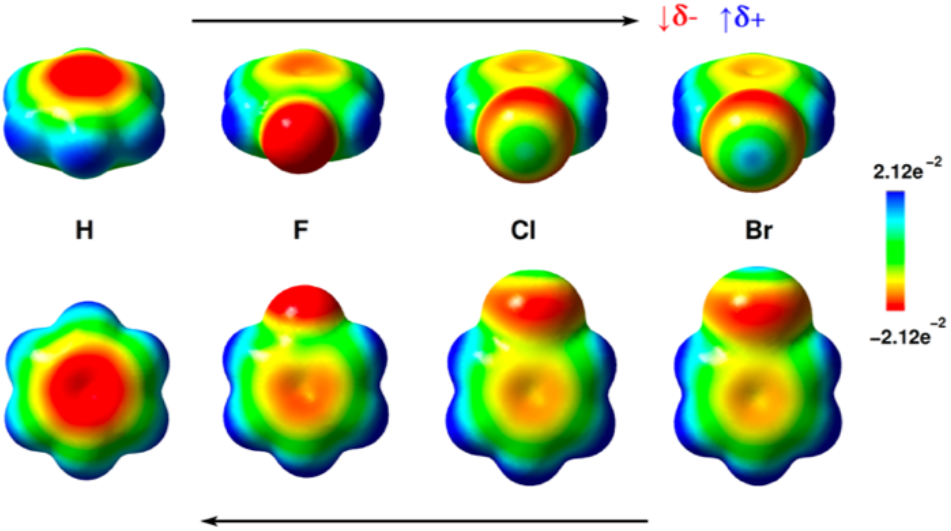
Electrostatic Surface Potential (ESP) maps (left to right) of benzene, fluorobenzene, chlorobenzene and bromobenzene at the 10^−3^ ea^-3^ isodensity surface. The color scale of the surface potential ranges from -0.0212 au (red) to 0.0212 au (blue). The upper black arrow indicates the increase in the sigma-hole strength of the halogens F < Cl <Br. The red arrow on the top right indicates the decrease of the electron rich region *δ*^−^ on the sides of the C–X bond in going from F to Cl to Br, and the blue arrow indicates the increase of the electron deficient region *δ*^+^ along the C–X bond from F to Cl to Br (X being the halogen). The lower black arrow indicates the increase of the negative electrostatic potential of the phenyl ring when passing from lager halogens to smaller to no halogens at all (Br<Cl<F<H).

### Alchemical Simulations of Halogenated Insulin Analogs Predict Significant Changes in Thermodynamic Stability

The simulations started from molecule 1 of the T_6_ zinc insulin hexamer (PDB ID 4INS refined at a resolution of 1.5 Å)^20^ using the “all atom” protein force field (CHARMM22) with provisions for multipolar interactions and periodic boundary conditions. The force field included a correction for α-helical bias by (Φ,ψ) CMAP potential (see SI for more details). Energy minimization yielded an essentially identical structure; root-mean-square differences (RMSD) for residues B3-B28 and A2-A20 were 0.6 Å (main-chain) and 1.8 Å (side chain). Halogen-induced electrostatic dipole moments of halobenzenes (Fluorobenzene, chlorobenzene and bromobenzene) exhibited relative strengths F (1.21 D) > Cl (0.65 D) > Br (0.12 D). These values derived from electronic structure calculations.^28-30^ Respective alchemical mutation of aromatic hydrogens in the nine cases studied (2-F through 6-F in the first set of simulations; 2-Cl, 6-Cl, 2-Br and 6-Br in the second) was well tolerated without transmitted structural perturbations. Deviations in B24 side-chain dihedral angles were in each case <10°. Because of subtle asymmetries within the crystallographic dimer,^20^ simulations were repeated beginning with molecule 2; similar results were obtained (*i*.*e*., within statistical error of the computation).

Alchemical simulations (see Figure S1a in Supplementary Information (SI)) provide information about stability changes (enthalpic and entropic) upon mutation. First, the focus was on fluorous substitutions. The free energies were computed using thermodynamic integration (TI), where a scaling parameter λ is applied and switches between initial (λ=0, state A) and final (λ=1, state B) states by gradually damping all nonbonded interactions (see SI section I for more details). Estimates of ΔΔ*G* obtained from free energy simulations are summarized in Table 1; “negative/positive” values represent a “gain in/loss of” stability relative to WT upon halogenic modification. Most analogs were predicted to be more stable than the unmodified protein; stabilization was mostly marked at position 2 followed by position 6, which is related to position 2 by a 180° rotation of *χ*_2_ (“ring flip”) (see Figure S1b in SI). The average gain in stability of 2-F-Phe^B24^-insulin (ΔΔ*G* = -1.42±0.38 kcal/mol relative to WT) was predicted to be twofold larger than that of 6-F-Phe^B24^-insulin (ΔΔ*G* = -0.63±0.38 kcal/mol relative to WT) (with a 95% confidence interval of 1.5-4 fold). Unlike the effects of an *ortho*-fluoro substitution (for which position 2 is preferred over position 6), ring-flip-related *meta* isomers (3-F-Phe^B24^ and 5-F-Phe^B24^) exhibited similar predicted stabilities (ΔΔG = -0.38 ±0.21 and -0.45±0.21 kcal/mol for 3-*meta* and 5-*meta*, respectively). At the *para* position (4-F-Phe^B24^-insulin) the null hypothesis (unchanged stability) could not be excluded.

**Table 1.** Stability free energy differences (ΔΔ*G*_Stab_) of halogenated Phe^B24^ insulin analogs. As a baseline validation, control alchemical “mutations” of a hydrogen to a hydrogen in Phe^B24^ in the models yielded ΔΔ*G* ≈ 0.

| $\Delta\Delta G_{\text{stab}}$<br>(kcal/mol) | | 2-*ortho* | 3-*meta* | 4-*para* | 5-*meta* | 6-*ortho* |
| --- | --- | --- | --- | --- | --- | --- |
| F-Phe<sup>B24</sup> | Theor. | $-1.42 \pm 0.38$ | $-0.38 \pm 0.21$ | $-0.21 \pm 0.21$ | $-0.45 \pm 0.21$ | $-0.63 \pm 0.38$ |
| | Exptl. | $-1.0 \pm 0.16$ | $-0.2 \pm 0.16$ | $0.05 \pm 0.16$ | - | - |
| Cl-Phe<sup>B24</sup> | Theor. | $-1.38 \pm 0.42$ | - | - | - | $0.07 \pm 0.42$ |
| | Exptl. | $-1.0 \pm 0.12$ | - | - | - | - |
| Br-Phe<sup>B24</sup> | Theor. | $-1.09 \pm 0.52$ | - | - | - | $1.11 \pm 0.52$ |
| | Exptl. | $-0.8 \pm 0.11$ | - | - | - | - |

Given the favorable predicted effects for *ortho*-F-Phe^B24^, the simulations were extended to larger halogenic substitutions at *ortho* positions 2 and 6. Initial energy minimization suggested Cl and Br would be readily accommodated whereas iodine was too bulky in this position and so was not considered further. This differs from the situation for Tyr^B26^ for which an iodinated residue can be readily accommodated. Results for these four additional simulations are given in Table 1. A trend was observed wherein (i) F, Cl and Br substitutions at ring position 2 are each stabilizing whereas (ii) at ring position 6 only the fluorous substitution was stabilizing; larger halogens at the 6-position were destabilizing.

### The net changes in free energy result from a large number of small effects

To relate predicted changes in thermodynamic stability to potential conformational changes, ten additional MD simulations each 10 ns in length were carried out: the WT monomer; F-Phe^B24^ in positions 2, 3, 4, 5 and 6; and the Cl-Phe^B24^ and Br-Phe^B24^ variants in the two *ortho* positions (2 and 6). The simulations were analyzed to evaluate possible structural origins of the predicted changes in stabilities in terms of (i) flexibility of the B-chain (B24 to B29) (Figures S2-S4 in SI), (ii) new local interactions (Figures 3 and S5 in SI) and conformational rearrangements (Figures 4a and S6-S7 in SI) and (iii) individual energy contributions (van der Waals and electrostatic interactions; see Figures S8, S9 and Tables S1, S2 in the SI). This analysis suggests: (i) B-chain flexibility (B24-B29) depends on the type and position of the halogen; (ii) each halogenation induces distinguishable local interactions, which drive conformational rearrangements; and (iii) imperfect enthalpy-entropy compensation (EEC) rationalizes stabilization of ring position 2 over 6.

**Figure 3.**
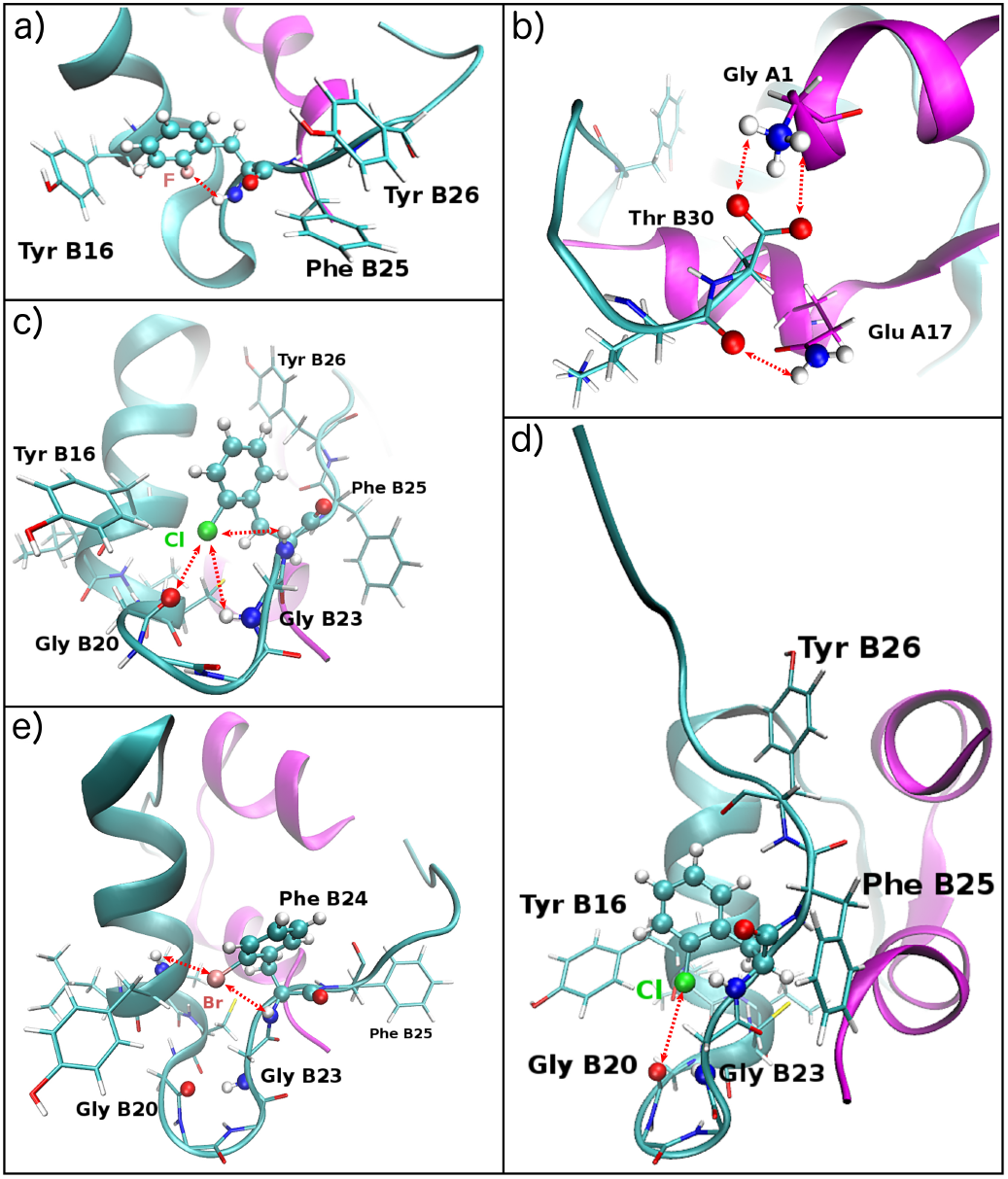
Structures of halogenated 2-X-Phe^B24^ insulin monomers. **a**. Perpendicular *π*-stacking of 2-F-Phe^B24^ and Tyr^B26^ induced by the interaction of F with the –NH backbone atom of 2-F-Phe^B24^. **b**. Interactions between Thr^B30^ and residues Gly 1 and Glu 17 of chain A caused by the “2-F-Phe^B24^”-induced conformational change locked state. The B-chain C-terminus is locked between the 2 helices of chain-A. **c**. Interactions of Chlorine with the backbone atoms of the B-chain crevice. The figure illustrates the formation of a sigma-hole interaction between the *δ*^+^ region of Cl and the O backbone atom of Gly^B20^. **d**. Bending of the B-chain C-terminus away from A-chain due to the halogen bond formation. **e**. Interactions of bromine with the –NH backbone atom of 2-Br-Phe^B24^ and Tyr^B16^, respectively; Br bridges between the *β*–strand and the B-chain helix. Red dashed arrows indicate the hydrogen/halogen-bonds.

**Figure 4.**
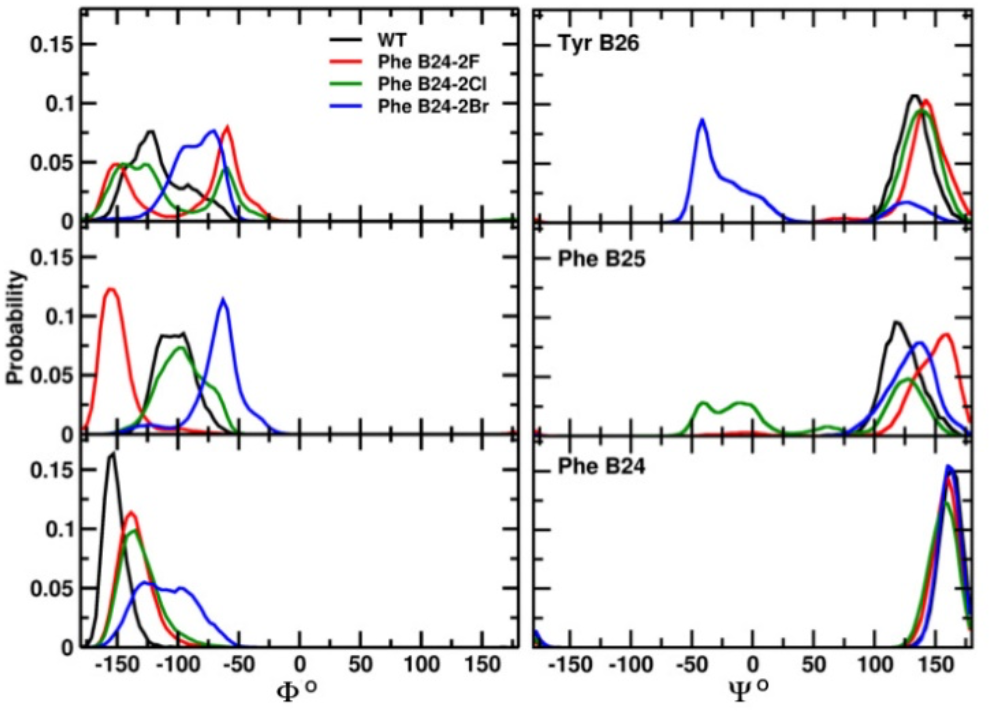
Backbone dihedral angle distributions of Phe^B24^, Phe^B25^ and Tyr^B26^ as a function of the halogen introduced in ortho-2 to Phe^B24^.

### Flexibility of B-chain C-terminal segment (B24-B29) depends on specific B24 modification

The displacement of the C-terminal segment of the B chain residues with respect to Gly^A1^ is of interest due to its immediate relevance to the mechanism of receptor binding and was first investigated for the fluorinated Phe^B24^ monomers. The evolution of the distance between the N-backbone atom of residue A1 and the C=O backbone atom of residue B30 was monitored (“*d*_A1-B30_”; Figure S2a). For the WT monomer the distance progressively increased from around 10 Å to 15 Å during the MD simulation compared with the fluorinated proteins which exhibited larger distance fluctuations ranging from 10 to 22 Å. Nevertheless, in some cases (4-F-Phe^B24^, 5-F-Phe^B24^ and especially for 2-F-Phe^B24^) the distance also decreased to 2.5-3 Å in the course of the trajectories. For 2-F-Phe^B24^ variant (lower panel in Figure S2a, black line) *d*_A1-B30_ decreased from ∼12.5 to 2.5 Å during the first 3 ns and then maintained this closed conformation for the majority of the simulation. Figure S2b shows RMSD values relative to the starting X-ray conformation (residues B18-B30). α-Helical conformations were stable in each MD simulation. The RMSD from the starting conformation is ∼1 Å (data not shown). In contrast, the N-terminal and the C-terminal ends of B-chain were flexible. For WT insulin, RMSD values of the B-chain C-terminal segment increased from ∼1.0 to 1.5 Å during the first 2 ns of the MD simulation and then stabilized around 2.5 Å. The fluoro-Phe^B24^ analogs each exhibited higher RMSD values, beginning at 2 Å and increasing to 4 Å (after 1 ns for F in *para*, in *meta* and in *ortho*-6 positions or after 4 ns for F in *ortho*-2; Figure S2b). Superposition of frames sampled from these trajectories (see Figure S2c in relation to the reference PDB monomer 4INS; red) enabled monitoring of conformational changes of the B-chain C-terminal segment. These structures reflect the above trends in *d*_A1-B30_ values (Figure S2a), including the “locked” conformation of the 2-F-Phe^B24^ analog. Profound differences were seen in the simulations of corresponding 2-Cl-Phe^B24^ and 2-Br-Phe^B24^ analogs (Figure S3). Whereas 2-F led to occlusion of the A1-A3 region, this receptor-binding surface was in part re-opened by 2-Cl-Phe^B24^; 2-Br-Phe^B24^ was associated with an entirely new conformation. Similar profound differences were also observed for *ortho*-6 halogenated analogs (Figure S4). These effects are discussed below in relation to backbone dihedral-angle distributions.

### B24 modifications introduce new local interactions

The above MD simulations suggested that the B-chain C-terminal segment can differ in flexibility depending on the modification. To explore the origins of these effects, the backbone Φ− and ψ−dihedral angle distributions *P*(Φ) and *P*(ψ) for X-Phe^B24^, Phe^B25^ and Tyr^B26^ were considered next. The distribution functions for X-Phe^B24^ variants are shown in Figures S6-S7 and Figure 4 relative to WT. Figure S6 presents *P*(Φ) and *P*(ψ) of F-Phe^B24^, Phe^B25^ and Tyr^B26^ for the five fluorinated analogs. Whereas in each case B24 *P*(ψ), B26 *P*(ψ) and B24 *P*(Φ) were similar to WT, marked differences were observed in B26 *P*(Φ) and in both B25 *P*(ψ) *P*(Φ) specifically for2- and 6-F.

The B-chain bent away from its WT orientation at Tyr^B26^ (Figure S6, top panel, solid red line); this was apparently due to an intra-residue interaction between the F and the B24 backbone N-H (Figure 3a). This interaction required a change in B24 Φ angle from -150° to - 125°, in turn associated with a change in B25 Φ from -100° to -160° (middle panel) and leading to two B26 conformational substates, *open* and *closed* (the former with Φ = -150° and latter at Φ = -50°; *versus* -125° for WT). The acute Φ angle (−50°) oriented the C-terminal B-chain segment (B26-B30) toward the A chain. In this closed conformation Thr^B30^ engages in interactions with both the α-amino group of Gly^A1^ and the δ–NH_2_ of Glu^A17^ (Figure 3b). Although B24 main-chain dihedral angles were not affected (Figure S6, bottom panel, dashed red line), Φ for B25 decreased by 50° relative to WT (middle panel), and a new signature in *P*(ψ) appeared at 10° with the same probability as 125°. This was associated with a shift in B26 *P*(ψ) from -125° to -60° (upper panel). This concerted set of conformational changes presumably reflected electrostatic interactions between the 6-F and the amide–NH main-chain atoms of Phe^B25^ and Tyr^B26^ (Figure S5).

Figure 4 focuses on the three modifications at position 2 (X=F, Cl, Br). Comparison of the *P*(Φ) and *P*(ψ) distributions of Phe^B24^, Phe^B25^ and Tyr^B26^ highlighted local changes in the orientation and interactions of nearby side chains. Like 2-F-, 2-Br-was associated with a shift the *P*(Φ) peak from -150° (WT) to -125°, whereas 2-Cl-was associated with a wider distribution ranging from -125° to -80°. Distinct patterns of transmitted conformational changes were observed. Whereas 2-Cl-didn’t shift from the B25 *P*(Φ) WT distribution, 2-F- and 2-Br-were associated with respective shifts in B25 *P*(Φ) peak to -150°and to -60°. 2-Cl-, the emergence of a new *P*(ψ), compared to the WT (Δ 125°), ranging from -50° to 0°. For Tyr^B26^, 2-Cl-invokes similar modification to *P*(Φ) as 2-F-, but with different probabilities, while 2-Br-leads to the appearance of a wide distribution ranging from -100° to -60°. Br also induces a shift of the Tyr^B26^ *P*(ψ) peak from Δ 140° (for WT, 2-F- and 2-Cl-) to -40°.

The dynamic behavior of the B-chain C-terminal segment in the 2-Cl-Phe^B24^ mutant can be related to the Cl atom interacting with positive and negative sites of the protein via both its electron rich δ^-^ and electron deficient δ^+^ regions (Figure 3c). The δ^+^ region forms a σ-hole bond with the carbonyl–O backbone of Gly^B20^ and the δ^-^ region interacts with the amide–NH backbone atoms of Gly^B23^ and Phe^B24^, between which it alternates. Due to the Cl--O α-hole bond, the Phe^B24^ residue is pulled towards the U-turn which explains the B-chain bending away from A-chain (Figure 3d). For the 2-Br-Phe^B24^ mutant, the large vdW radius of Br allows the formation of an H-bond with the amide–NH backbone atom of Phe^B24^ and Tyr^B16^, respectively (Figure 3). Br bridges between the Δ-strand and the α-helix of chain-B bringing them closer to each other at the Tyr^B16^ – Phe^B24^ level, which causes the U-turn to tighten. The backbone carbonyl-O of Gly^B20^ is then pushed to larger distances from Br and no σ-hole bond is formed.

Figure S7 compares Phe^B24^ halogenation in the *ortho*-6 position relative to WT; the *P*(Φ) peak of Phe^B24^ shifts from -150° to -135° and -115° when passing to Cl and Br, respectively. In this position, the halogens are exposed to the backbone carbonyl-O of Phe^B24^ and thus the decrease of *P*(Φ) when passing from F to Cl to Br is dependent on the size of the halogen introduced. The *P*(ψ) of Phe^B24^ shows in the case of Br the emergence of a new state at 125°. For Phe^B25^, 6-F- and 6-Cl-shifts the *P*(Φ) peak from -100° to -135° and -140° respectively, while 6-Br-broadens the angular distribution peak from -100° to - 75°, and shifts the *P*(ψ) from 75° to 15° and -45°. The effect of the 6X-halogenations on Tyr^B26^ affects only *P*(Φ), where F, Cl and Br shift the peak from -125° to -60° but with a lower probability in the case of Br. The 6-Br-mutation also shows the emergence of a second peak at -145° with a higher probability than the peak at -60°. The dihedral angular variations induced by the 6-Cl-mutation can be explained by the halogen bond formation between the latter and the amide–NH backbone atoms of Phe^B25^ and Tyr^B26^, as in 6-F-(Figure S5a,b). However, the accommodation of the larger vdW radius of Cl, compared to F, induces the emergence of new *P*(Φ) and *P*(ψ) for Phe^B25^. The largest *P*(Φ) and *P*(ψ) emerge from the 6-Br-halogenation. In this case, there is a σ-hole bond formation between the δ^+^ region of Br and the carbonyl-O backbone of Phe^B24^ in a first step (Figure S5c), and in a second step the B24 ring flips and Br interacts with the amide–NH backbone atom via its δ^-^ region (Figure S5d); this observation explains the emergence of two different peaks in the dihedral angle distributions.

### Enthalpy-Entropy Compensation Explains Stabilization of 2X versus 6X

Interaction energies (*ΔE*_*int*_) were analyzed for halogen substituents at positions 2 and 6 (Figure S8 and S9, respectively), as calculated from the above 10 ns MD simulations. The analyses focused on two sets of interactions between (i) the B24 side chain and the A-chain, and (ii) the B24 side chain and the B-chain. Non-bonded energy contributions were decomposed into an electrostatic *(ΔE*_*elec*_; red lines) and a van der Waals *(ΔE*_*vdW*_; green lines) component.

#### (i) Position 2

Relative to WT, halogen atoms at position 2 generally increased interaction energies between the modified side chain and the A-chain (Figure S8a). In 2-F-Phe^B24^, *ΔE*_*int*_ increased from -2 (WT) to *ca*. -4 kcal/mol (Figure S8a); the interaction energy with the B-chain decreased by ∼ 5 kcal/mol (Figure S8b). In each case the total energy was mirrored by its electrostatic contribution *ΔE*_*elec*_ (red lines). Thus, 2-F increased (decreased) the interaction energy with A-chain (B-chain) residues. This trend was associated with the movement of the B-chain C-terminus towards the N-terminal A-chain α-helix and away from the central B-chain α-helix (Figure S3c).

For 2-Cl-Phe^B24^ the increased *ΔE*_*int*_ was driven almost entirely by increased interactions with the B-chain (∼ 5 kcal/mol; Figure S8b) whereas *ΔE*_*int*_ with the A-chain was not significant (Figure S8a) in accordance with structural trends opposite to those induced by 2-F (Figure S3c). A striking implication is that 2-F and 2-Br can each achieve net enthalpic stabilization via distinct atomic mechanisms. The pattern of interaction energies observed in 2-Br-Phe^B24^ was similar to 2-F. The larger structural excursions in this case (Figure S3c) were associated with larger fluctuations in interaction energies. 2-Br enhanced *ΔE*_*int*_ between B24 and A-chain residues after 1 ns of MD simulation (ranging between -2 and -8 kcal/mol; Figure S8a) and *ΔE*_*int*_ between B24 and B-chain residues (ranging from -7.5 to 7.5 kcal/mol; Figure S8b). This trend was associated with distinct structural changes in the C-terminal B-chain segment, arising from Φ and ψ angular variations of the base residues Phe^B24^, Phe^B25^ and Tyr^B26^ (Figure 4). These changes enabled the C-terminal B-chain residues to engage with both the N-terminal A-chain α-helix and central B-chain α-helix (Figure S3c).

Relative to 2-F, the total energy profiles in the larger halogens were primarily driven by its *ΔE*_*vdW*_ contribution. The more substantial fluctuations of *ΔE*_*int*_ in the 2-Cl and 2-Br simulations were each associated with greater flexibility of the B-chain C-terminal segment (Figure S3).

#### (ii) Position 6

Electrostatic terms made a leading contribution at position 6; *ΔE*_*vdW*_ terms were similar among the halogens (Figure S9b). The halogen substituents at position 6 induced large energy fluctuations associated with flexibility of the C-terminal B-chain segment (Figure S4). This trend was more pronounced than in the 2-X simulations as observed by RMSD values and *d*_A1-B30_ evaluations (Figure S4a,b)). These findings predict increased changes in configurational entropy relative to the 2-X simulations.

In 6-F-Phe^B24^, the C-terminal B-chain segment alternated between two states, one leaning toward the A-chain and the other leaning away (Figure S4c). These two states underlay the increased *ΔE*_*int*_ between 6-F-Phe^B24^ and residues in both chains (Figure S9a,b). For 6-Cl-Phe^B24^ *ΔE*_*int*_ was increased between 6-Cl-Phe^B24^ and the-A chain but decreased between Phe^B24^ and the B-chain. This trend was associated with preferred orientation of the C-terminal B-chain segment toward the A-chain. Changes in *ΔE*_*int*_ terms for 6-Br-Phe^B24^ and structural displacements of the B-chain C-terminal segment were opposite to that of 6-Cl (Figure S9a,b and Figure S4c).

Residues in insulin affected by the B24 modifications at positions 2 or 6 were identified (Tables S1 and S2, respectively). These tables provide a comparison of vdW (Δ*E*_vdW_) and electrostatic (Δ*E*_elec_) energy fluctuation differences of backbone (bb) and side-chain (sc) atoms relative to WT, as calculated from the 10 ns MD simulation. In addition to the modified residue itself (B24), major enthalpic fluctuations were found to involve residues Gly^A1^, Glu^A4^, Glu^A17^, Tyr^A19^, Cys^A20^, Gln^A21^, Val^B12^, Leu^B15^, Tyr^B16^, Cys^B19^, Gly^B20^, Arg^B22^, Gly^B23^, Phe^B25^, and Tyr^B26^. Most of these residues are not in direct contact with Phe^B24^.

These results are in accordance with the observed interactions (Figure 3 and Figure S5) and with the above analysis of segmental conformational changes (Figure S2-S7 and Figure 4). Favorable changes in enthalpy were dipole dependent in the series (F > Cl > Br). Predicted effects are more marked at position 2 (−9.58 > -8.94> -8.81 kcal/mol) than at position 6 (−7.08 > -3.15 > -2.67 kcal/mol). As indicated by the free-energy calculations in Table 1, changes in entropy following modification at position 2 cannot compensate this highly increased enthalpy, whereas at position 6 the enthalpic terms are substantially offset by compensating entropic terms. These simulations thus predict that *ortho*-X-Phe^B24^ insulin analogs will predominantly occupy the 2 position in a conformational equilibrium related by χ_2_ angle.

Together, the above simulations suggest that net changes in free energy result rather from a large number of small effects than one single dominant intermolecular interaction. Surprisingly, the stabilizing effects at position 2 had different microscopic origins depending on the type of halogen. In each case EEC occurred in association with dynamic changes in the modified B-chain. As expected, predicted changes in ΔΔ*G* are generally smaller than components ΔΔ*E* (as a surrogate for ΔΔ*H*) and *T*ΔΔ*S* (Table 2). Relative strengths of the halogen-induced dipoles (F > Cl > Br) inversely correlate with extent of EEC. Hence, changes in entropy cannot mitigate favorable changes in enthalpy. Conversely, corresponding changes in enthalpy at position 6 can not compensate for unfavourable changes in entropy. As a general feature of these systems, gains in solvation entropy play a dominant role (arising from the displacement of the water molecules; some of these water molecules play an important role in the stability of insulin dimers)^31^ in lowering free energies.^32, 33^ The simulations are thus consistent with the perspective that lock-and-key binding is an entropy-dominated process.^34^ Predicted decreases in total Gibbs free energies reflect a delicate balance of opposing effects. This outcome is remarkable given that EEC often confounds efforts to engineer more stable proteins.

**Table 2.** Entropy-enthalpy compensation. Free energies (ΔΔ*G*) were decomposed into respective enthalpic (ΔΔ*H*) and entropic (ΔΔ*S*) contributions in the monomeric protein analogs (ΔΔ*G* data taken from Table 1). ΔΔ*H* was calculated from 10 ns of MD simulations (ΔΔ*H* data taken from Tables S1 and S2 for 2-*ortho* and 6-*ortho* positions, respectively). ΔΔ*S* was calculated according to ΔΔ*G* = ΔΔ*H* – *T* ΔΔ*S* at *T* = 300 K; units are in kcal/mol. ΔΔ*H* and ΔΔ*S* are averages over 10 ns simulations.

| halogen | position 2-*ortho* |  |  | position 6-*ortho* |  |  |
| --- | --- | --- | --- | --- | --- | --- |
| | $\Delta\Delta G$ | $\Delta\Delta H$ | $-T\Delta\Delta S$ | $\Delta\Delta G$ | $\Delta\Delta H$ | $-T\Delta\Delta S$ |
| F | -1.42 | -9.58 | 8.16 | -0.63 | -7.08 | 6.45 |
| Cl | -1.38 | -8.94 | 7.56 | 0.07 | -3.15 | 3.22 |
| Br | -1.09 | -8.81 | 7.72 | 1.11 | -2.67 | 3.78 |

### Evaluation of Corresponding Insulin Analogs Confirms Theoretical Trends

To experimentally test these predictions, halogenic substitutions were introduced within an engineered insulin monomer of native activity.^26^ This engineered insulin monomer, designated as KP-insulin (or insulin *lispro*), the parent analog of the present halogenic insulins contains two substitutions in the C-terminal segment of the B-chain that impede dimerization (Pro^B28^→Lys and Lys^B29^→Pro).^35^ Use of this template circumvented confounding effects of self-assembly on stability assays. The structure of KP-insulin in solution closely resembles a crystallographic T-state protomer.^36^

Five analogs of KP-insulin were prepared by semi-synthesis.^37, 38^ The three fluorous analogs were 2/6-F-Phe^B24^ (collectively designated *ortho*-F-Phe^B24^-KP-insulin), 3/5-F-Phe^B24^ (*meta*-F-Phe^B24^-KP-insulin), and 4-F-Phe^B24^ (*para*). In addition, *ortho*-Cl-Phe^B24^-KP-insulin and *ortho*-Br-Phe^B24^-KP-insulin were prepared to evaluate halogen-specific effects. Far-ultraviolet circular dichroism (CD) spectra of these analogs are essentially identical to that of KP-insulin (Figure S10 in SI).

Thermodynamic stabilities, as inferred from CD-detected chemical denaturation,^26^ are given in Table S3. Strikingly, *ortho*-F-Phe^B24^-KP-insulin exhibited markedly increased stability (ΔΔ*G*_u_ = -1.0±0.2 kcal/mol relative to KP-insulin) whereas *meta*-F-Phe^B24^ exhibited little gain in stability (ΔΔ*G* = -0.2±0.2 kcal/mol; the *para*-Phe^B24^ analog was similar in stability (ΔΔ*G*_u_ = 0.05±0.2 kcal/mol; Figure 5a and Table S3, SI). The fluorous substitutions also have reciprocal effects on *m-*values (Table S3), which correlate with the extent of change in exposure of hydrophobic surfaces upon denaturation.^39^ The larger two halogens at the *ortho* position were also stabilizing but less so than fluorine (Figure 5b).

**Figure 5.**
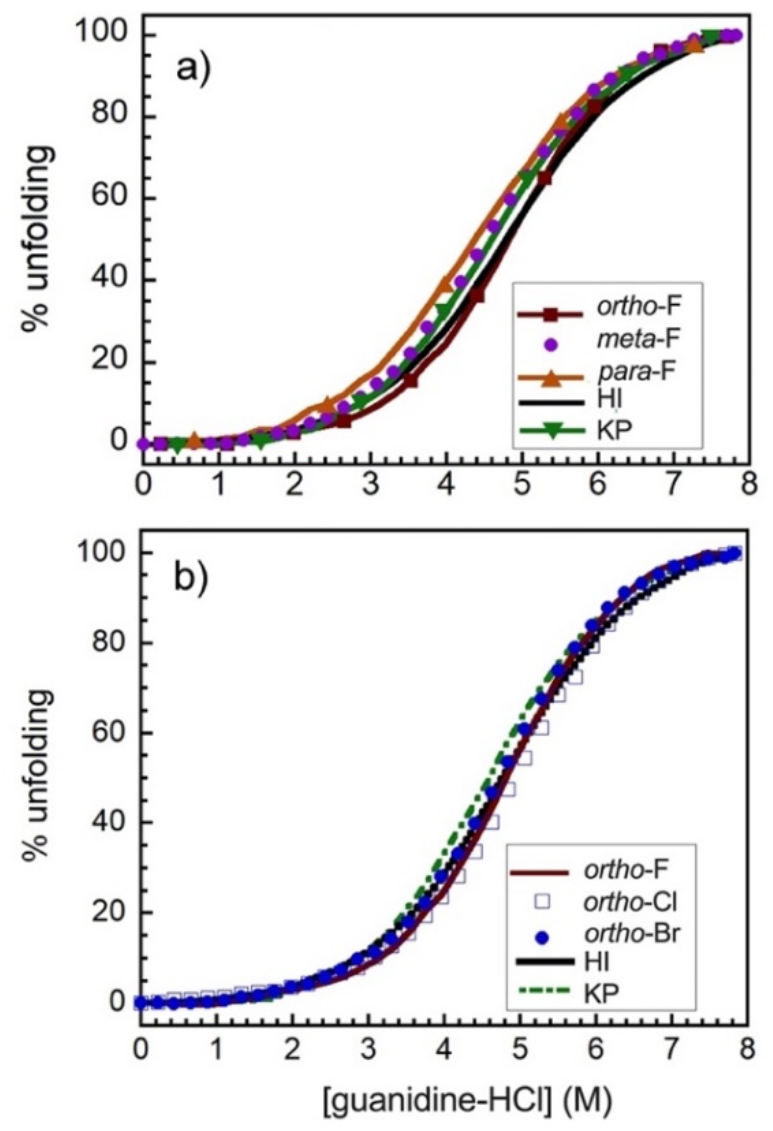
Thermodynamic stability of insulin analogs. Guanidine denaturation assays of insulin analogs **a**. substitution of fluorine atom at *ortho, meta* and *para* position of Phe^B24^ and **b**. substitution of various halogens (F, Cl and Br) at *ortho* position along with controls human insulin (HI) and insulin lispro (KP), monitored by ellipticity at 222 nm. The resulting stability values are given in Table S3 in SI.

### B24-related pocket couldn’t accommodate the dipole within the hormone-receptor complex

Affinities of the halogen-modified insulin analogs for the lectin-purified human insulin receptor (isoform A as immobilized in a competitive-displacement plate assay) were generally lower than that of KP-insulin (Figure 6 and Table S3). At the *ortho* position the affinities followed the rank order WT (strongest) < Br ∼ Cl < F (weakest), presumably reflecting relative electrostatic perturbations within the B24-binding pocket at the hormone-receptor interface.^21^

**Figure 6.**
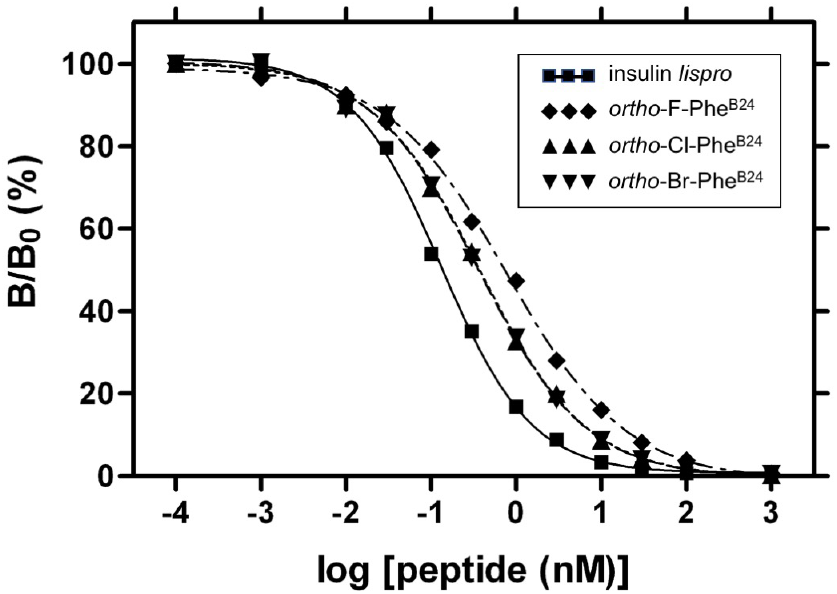
Receptor binding assay for insulin receptor (IR) for halogen analogs (*ortho*-F-Phe^B24^-KP-insulin ♦, *ortho*-Cl-Phe^B24^-KP-insulin ▴ and *ortho*-Br-Phe^B24^-KP-insulin ▾) compared to insulin lispro (▪). Affinities of the *ortho*-Cl and *ortho*-Br analogs were indistinguishable within experimental error.

To rationalize such affinity changes, hormone-IR interaction energies were evaluated from 1 ns of MD (see SI for the detailed analysis and Figure 7; For a non-halogen control provided by CH_3_ see Figure S11). This analysis highlighted the importance of electrostatic interactions: the larger the dipole (F > Cl > Br > CH_3_ = 0) the worse is the binding (F < Cl < Br < CH_3_). This perturbation was in part mitigated by favorable vdW contributions, more effectively in the two larger halogens. Entropic changes were consistently unfavorable.

**Figure 7.**
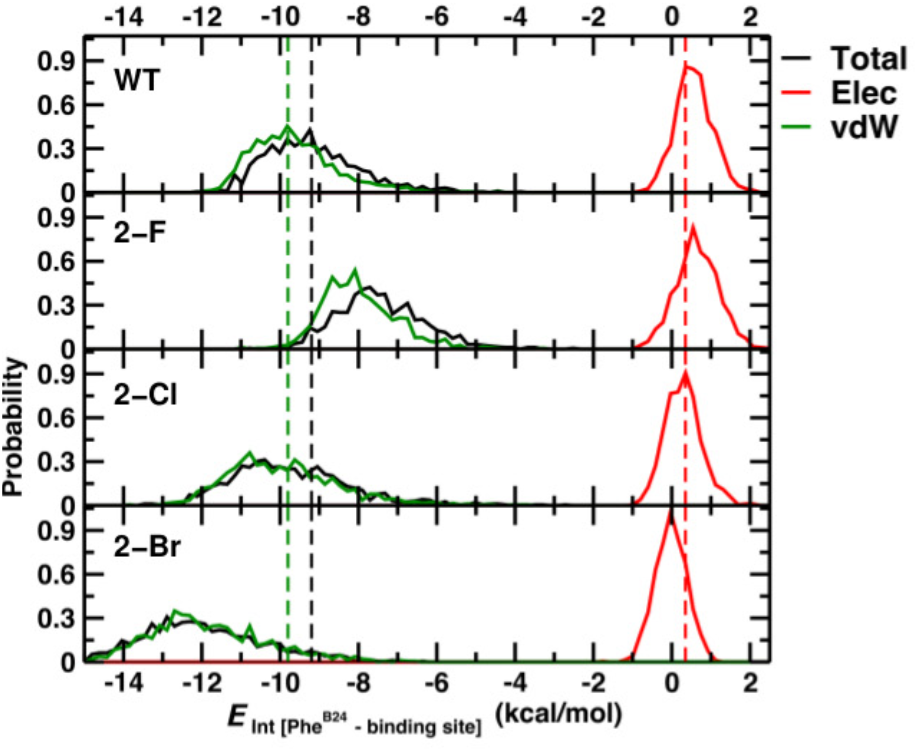
Probability distributions for the interaction energy between INS monomer and micro-receptor (μIR) from 1 ns of MD simulations for the 2-X halo-aromatic substitutions. Dashed lines represent averaged energy values of the WT-INS bound to the μIR fragments, as averaged over the 1 ns MD simulation. A complete set of results for ortho-2 and ortho-6, including CH_3_ as reference, is provided in Figure S11 (SI).

### Fluorination at the 2-Position Decreases Activity of the Insulin Monomer Towards the Receptor Through Self-Interaction

The unfavourable effect of 2-F-Phe^B24^ on IR binding was reinforced by self-interactions in the free analog simulation. The latter exhibited two conformational states (one at Φ = -150° and another at Φ = -50°) in *P*(Φ) for residue Tyr^B26^, compared to -125° for the WT (Figure 4). These states correspond to: (i) an open state wherein the B-chain C-terminal segment detached from the A chain to its nonpolar receptor-binding residues (Ile^A2^ and Val^A3^), and (ii) a closed state wherein the B-chain C-terminal segment is “locked” between the A-chain α-helices, effectively masking the A2 and A3 side chains (Figure S2 and Figures 3a,b). The higher the occupancy of the locked state, the less active would be the analog. Whereas MD simulations indicated a preference of the 2-F-Phe^B24^ analog for the locked state, 2-Cl-Phe^B24^ and 2-Br-Phe^B24^ modifications favoured the open states (Figure 4).

### Ortho-modified Insulin Analogs Retain Biological Activity in Mammalian Cells and in Rat Model of Diabetes Mellitus

Despite the above changes in IR affinity, biological activities of the analogs were found to be similar in male Lewis rats as rendered diabetic by pancreatic β-cell toxin streptozotocin.^26^ Studies were conducted following intravenous (i.v.) bolus injection (Figure 8; also see Figure S12 in SI). Such maintenance of activity in vivo has been observed in previous studies of low-affinity analogs and ascribed to delayed receptor-mediated clearance from the blood stream.^40-42^ Affinity-related reduction in insulin signaling could nonetheless be demonstrated in the case of *ortho*-F-Phe^B24^-KP in cell culture at 50 nM hormone concentration (see Figure S13 in SI)—assays not confounded by clearance. The robustness of insulin action *in vivo* to reduction in IR affinity (in the range 10-100%) has facilitated the design of clinical insulin analogs.^43^

**Figure 8.**
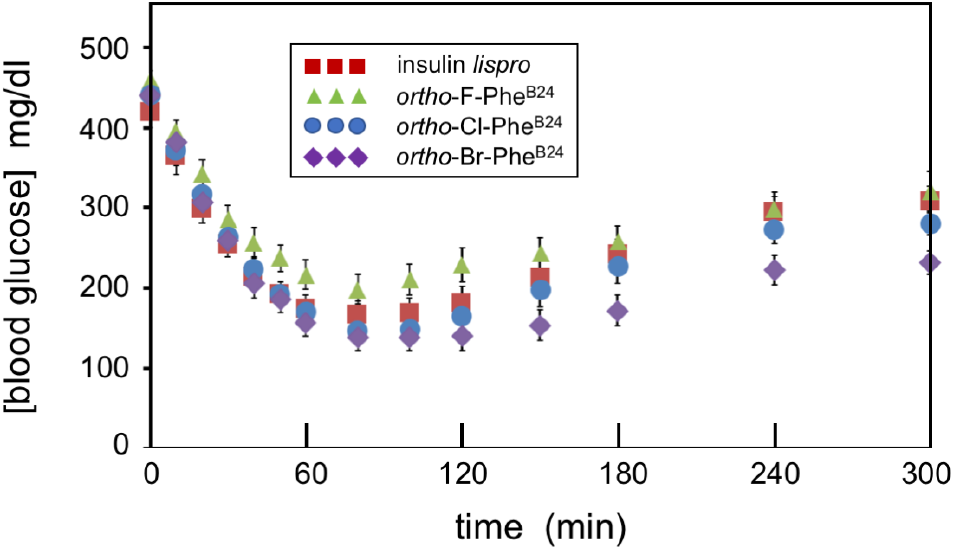
Rat studies of insulin analogues. Time course of [blood glucose] following i.v. injection. Insulin lispro 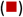; *ortho*-F-Phe^B24^-KP-insulin 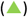; *ortho*-Cl-Phe^B24^-KP-insulin 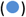; *ortho*-Br-Phe^B24^-KP-insulin 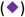. The doses of insulin analogs were 1.7 nmol per 300-gram rat (*n* = 6). For the diluent control, see Figure S12 in SI.

### Halogenic modifications of Phe^B24^ delay Onset of Protein Fibrillation

Whereas the focus so far has been on native-state stability to relative susceptibilities of insulin analogs to chemical degradation, ^1^ the shelf life of pharmaceutical formulations is also influenced by protein fibrillation.^44^ Lag times prior to onset of fibrillation depend not only on native-state stability, but also on state of self-assembly, conformational fluctuations leading to partial unfolding, and factors that modulate the formation or extension of an amyloidogenic nucleus (pH, temperature, ionic strength, excipients and protein concentration). Due to the complexity of this conformational transition, lag times do not correlate with standard free energy changes of unfolding.^45^ To evaluate the effect of halogenic substitutions of Phe^B24^ on lag times, two independent assays were undertaken.

The first employs an accelerated fibrillation protocol long-established in preclinical pharmaceutical pipelines.^46^ In this assay a 96-well plate is agitated at 960 rpm and 37°C with continuous fluorescence-based interrogation of each well, containing a protein solution and fibrillation probe thioflavin T (ThT). The results indicate that lag times under these conditions are prolonged in accordance with augmented thermodynamic stabilities (Figure 9A). The second assay instead mimics an insulin vial under real-world conditions: gentle movement (“sloshing”) of an insulin solution in a glass vial (∼60 rotations per minute at 37°C) in the presence of an air-liquid interface. In this slower assay aliquots were probed by ThT fluorescence, added subsequently as an indicator. Values are also reported in Supplemental Table S4 and S5. The results indicate that halogens in general delay fibrillation irrespective of effects on native-state stability (Figure 9B). We speculate that the latter protective effect reflects unfavourable alignment of the halogen atoms in an amyloidogenic nucleus whereas the accelerated assay probes native-state fluctuations prior to nucleus formation.

**Figure 9.**
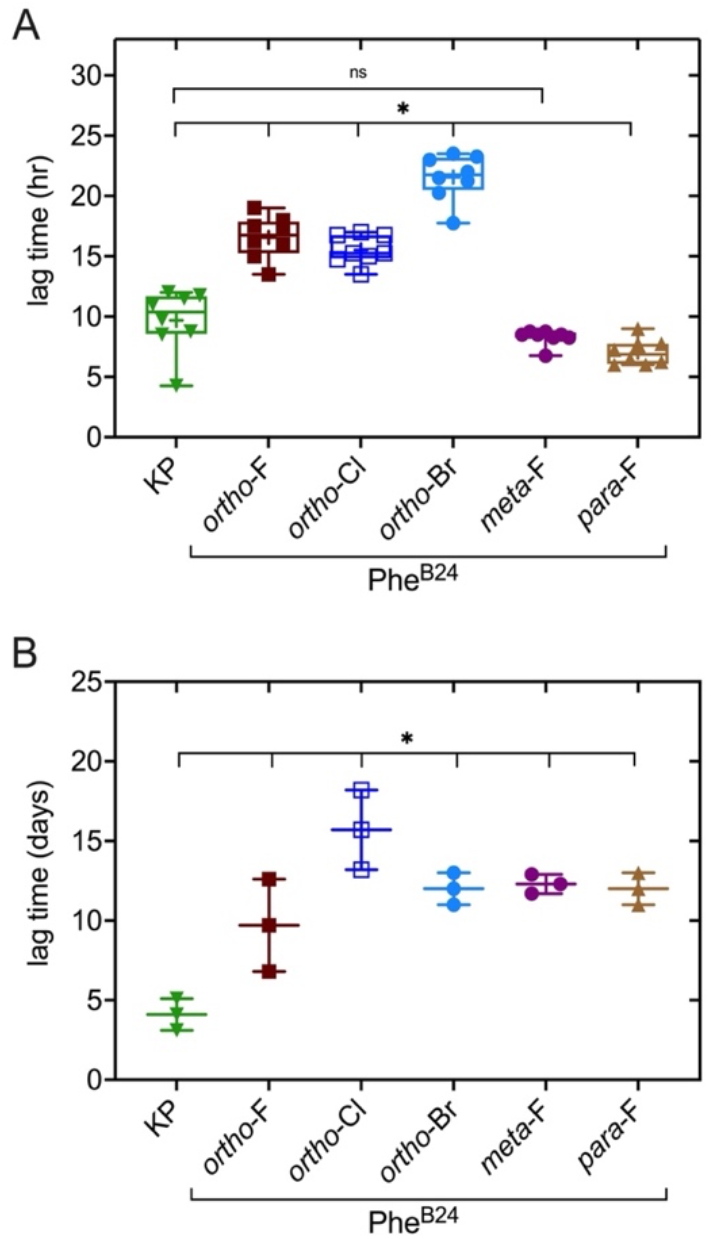
Analysis of fibrillation lag-times. Dot plot of lag times of various halogenated-Phe^B24^-KP analogs and KP-insulin by (A) accelerated protocol and (B) gentle rocking methods. Upper and lower borders of boxes respectively denote third and second quartiles; whiskers indicate 1.5-fold interquartile range from median. Mean values are shown by “+” sign. “ns” denotes not significant; asterisk (*) denotes significant and p-values <0.05.

## Concluding Remarks

Proteins are often stabilized by aromatic-aromatic interactions with geometric distance- and angular dependences.^18^ Such interactions arise from the asymmetric distribution of partial charges. Aromatic side chains may engage in a variety of weakly polar interactions, involving not only neighboring aromatic rings but also other sources of positive or negative electrostatic potential,^19^ such as peptide bonds and lone-pair electrons of disulfide bridges. The present study focused on a solvent-exposed crevice within a globular protein. We hypothesized that regiospecific distortion of π-orbitals and associated patterns of partial charges at ring edges lead to favorable or unfavorable changes in protein stability. Based on atomistic simulations it was predicted that nonlocal distortion of the aromatic system could act as an amplifier to enhance the per-halogen stabilization free energy.

Insulin contains three Phe residues (positions B1, B24, and B25). Because the exposed side chains of Phe^B1^ and Phe^B25^ are not well ordered in an insulin monomer,^47^ we focused on Phe^B24^. The amenability of insulin to semi-synthesis^37, 38^ facilitated incorporation of nonstandard side chains at this position. Conserved among vertebrate insulin-like factors, the aromatic ring of Phe^B24^ packs against (but not within) the hydrophobic core to stabilize the super-secondary structure of the B chain.^20^ In addition, Phe^B24^ lies at the receptor-binding surface where it participates in a change in conformation upon receptor binding.^21^ The marked and variable effects of halo-aromatic substitutions at B24 highlight its importance to both protein stability and receptor binding.

Strikingly, *ortho* modifications of Phe^B24^ are stabilizing whereas *meta*-F-Phe^B24^ and *para*-F-Phe^B24^ are neutral or destabilizing. Atomistic simulations highlighted favorable (or unfavorable) alignment of the induced electrostatic dipole moment of the modified aromatic ring within an asymmetric binding pocket. No single pairwise interaction was predominant. Although the halo-aromatic substitutions weakened binding to the isolated IR *in vitro* (in general accordance with the position of Phe^B24^ at the hormone-receptor interface^21, 38^), native-like activity was maintained in a rat model of diabetes mellitus.^48, 49^ In the future it will be of interest to determine the structures of these analogs and their IR-ectodomain complexes to ascertain the positions of the modified B24 rings and halogen atoms.

The developing world faces a challenge regarding storage, delivery, and use of drugs and vaccines.^50^ The requirement of a “cold chain” complicates the use of temperature-sensitive insulin formulations in regions of Africa and Asia lacking consistent access to electricity and refrigeration,^51^ a challenge likely to be deepened by an emerging pandemic of diabetes mellitus in the developing world.^52^ With continuing progress in methods for the manufacture of nonstandard proteins,^8^ we anticipate that halo-aryl substitutions may be employed (perhaps in combination with other stabilizing modifications) to design ultra-stable therapeutic products and vaccines.

## Conflicts of Interest

M. A. W. has equity in Thermalin, Inc. (Cleveland, OH) where he serves as Chief Innovation Officer; he has also been consultant to Merck Research Laboratories and DEKA Research & Development Corp. N. B. P. and F. I. -B. are consultants to Thermalin, Inc. and also have options, warrants or equity. L. W and J. W. has equity in Thermalin, Inc. F. I. -B. is a consultant to Sanofi and has received grants from Novo-Nordisk.

## Acknowledgements

We thank members of our respective laboratories for discussion; Q. X. Hua and Y. Yang for preliminary NMR studies; and Mads Krogsgaard Thomsenis (Novo Nordisk) for the gift of radio-labeled insulin. K. EH. and M. M. were supported by Swiss National Science Foundation Grant 200021-117810, National Competence Center for Research Molecular and Ultrafast Science and Technology (NCCR MUST). M. A. W. F. I. -B. and N. B. P. acknowledges the support of the U.S. National Institutes of Health (NIH R01 DK040949).

