## Supplemental Information for "Stabilization of a Protein by a Single Halogen-Based Aromatic Amplifier"

### Supplemental Information
for

### Stabilization of a Protein by a Single Halogen-Based Aromatic Amplifier

Krystel El Hage[a⊥^], Nelson B. Phillips[b^], Yen-Shan Chen[d], Balamurugan Dhayalan[d], Jonathan Whittaker[b], Kelley Carr[b], Linda Whittaker [b], Manijeh H. Phillips[b], Faramarz Ismail-Beigi[b,c], Markus Meuwly[a*], & Michael A. Weiss[b,d*]

[a.] *K. El Hage, M. Meuwly* Department of Chemistry, University of Basel, Klingelbergstrasse 80 CH-4056 Basel, Switzerland.*
[b.] *N.F. Phillips, J. Whittaker, K. Carr, L. Whittaker, M.H. Phillips, F. Ismail-Beigi, M. A. Weiss. Department of Biochemistry, Case Western Reserve University, Cleveland, Ohio 44106, USA.*
[c.] *F. Ismail-Beigi, Department of Medicine, Case Western Reserve University, Cleveland, Ohio 44106, USA.*
[d.] *Y.-S. Chen, B. Dhayalan, M. A. Weiss* Department of Biochemistry & Molecular Biology, Indiana University School of Medicine, Indianapolis, IN 46202, USA.*
⊥ Present address: SABNP, Univ. Evry, INSERM U1204, Université Paris-Saclay, 91025 Evry, France.
^ These authors contributed equally
*.

#### Purpose of Supplement

This Supplement is divided in two sections (Section I for Materials and Methods, and Section II for Ancillary Results). There are thirteen figures (Figure S1-S13), ten pertaining to simulations (Figure S1-S9 and S11) and three to experiments (Figure S10, S12 and S13). Also included are three tables (Tables S1-S3), two pertaining to simulations (Table S1-S2) and three to experiments (Table S3-S5). Please note that ring positions 2 and 6 of Phe are collectively designated *ortho*; ring positions 3 and 5, *meta*; and ring position 4, *para* (Figure S1b).

#### Table of Contents

Supplemental Methods……..2
Supplemental Simulations……..5
Supplemental References……..9
Figure S1……..10
Figure S2……..11
Figure S3……..12
Figure S4……..13
Figure S5……..14
Figure S6……..15
Figure S7……..16
Figure S8……..17
Figure S9……..18
Figure S10……..19
Figure S11……..20
Figure S12……..21
Figure S13……..22
Table S1……..23
Table S2……..24
Table S3……..25
Table S4……..25
Table S5……..25

### I. Supplemental Methods

**Molecular Dynamics Simulations.** Initial coordinates for the MD simulations were 2-Zn molecule 1 (1.5 Å; PDB 4INS).[1] MD simulations were performed with CHARMM[2] version c40a1 with the "all atom" protein force field (CHARMM22)[3] with provisions for multipolar interactions[4] and periodic boundary conditions were employed. The force field included a correction for α-helical bias as introduced into CHARMM22 by (Φ,Ψ) CMAP potential.[5] The insulin monomer was solvated in a 65.19 Å x 52.77 Å x 46.56 Å box of TIP3P water molecules[6] equilibrated at 300 K and 1 atm. Following 10000 steps of steepest-descent minimization, the systems were heated from 0-300 K for 15 ps and equilibrated for 50 ps, which was followed by 10 ns of production run. Equations of motion were propagated with the Verlet algorithm; the time step was $\Delta t$ = 1 fs. Bonds involving hydrogens were constrained with SHAKE.[7]

**Force Field Parametrization.** The bonded parameters for halogenated $Phe^{B24}$ were those from the CHARMM General Force Field (CGenFF).[8] Halogens bound in a molecule present on their outer side an electronically depleted region (the "σ-hole") that gives rise to a positive electrostatic potential along the carbon-halogen bond.[9] A multipolar (MTP) model, based on previously validated MTPs and Lennard Jones (LJ) parameters[4] was thus employed for the halogenated $Phe^{B24}$ rings (PhF, PhCl and PhBr) to improve the original point-charge (PC) force field in describing the σ-hole. The MTP model is a higher-order expansion of the electrostatic potential that allows a more realistic description of anisotropic electrostatic features. MTP parameters were derived from *ab initio* calculations at the MP2 level with the aug-cc-pVDZ basis set using GAUSSIAN09.[10] Electrostatic potentials ESPs were fit to MTP parameters in the first interaction belt. Non-bonded terms were scaled against MTP electrostatics for PhF, PhCl and PhBr, respectively, and re-parameterized by monitoring three thermodynamic quantities: pure-liquid densities, as well as heats of vaporization and hydration free energies in order to accurately reproduce experimental values.[4, 11] The experimental values of the three thermodynamic quantities were taken from the literature.[12]

**Free Energy Simulations.** For the free energy simulations, hybrid residues with dual topology were constructed for the modified side chains. Free energies were computed using thermodynamic integration (TI).[13] This protocol applies a scaling parameter λ and switches between initial (λ =0, state A) and final (λ =1, state B) states by gradually damping all nonbonded interactions. The Hamiltonian A→ B refers to the transformation between states A and B. Working in the slow-growth regime, the free energy is:

$$\Delta G_{A\to B} = \int_0^1 d\lambda \ \langle\frac{\partial\mathcal{H}}{\partial\lambda}\rangle_\lambda \ \approx \ \sum_i(\lambda_{i+1} - \lambda_i) \ \langle\frac{\partial\mathcal{H}}{\partial\lambda}\rangle_{\lambda m} \qquad (1)$$

where state A is the WT residue $Phe^{B24}$ and state B is the modified residue $Phe^{B24}$→X. The canonical average $\langle\,.\,\rangle_\lambda$ is performed over the phase space generated by $\mathcal{H}(\lambda)$, and $\lambda_m$= $(\lambda_{i+1} - \lambda_i)$. LJ and PC derivatives were obtained from the PERT module, using soft-core potentials for the LJ interactions.[13-14] Changes in free energy due to MTP electrostatics with coupling $\lambda_m$ were computed as described. [4, 11]

Simulations at constant pressure and temperature (*NpT*) were performed to compute the free energy at each $\lambda$ value using the Hoover heat-bath method with pressure coupling at $T$ = 298 K and $p$ = 1 atm; the masses of the temperature and pressure piston were *ca.* 20 % and 2 % of the system's mass, respectively. A friction coefficient of 50 $ps^{-1}$ was used. The interval $0 < \lambda < 1$ was broken up into 20 steps. At each step the system was re-equilibrated for 40 ps followed by 100 ps of dynamics. λ was changed from initial to final value using the slow-growth protocol, which allowed the system to re-equilibrate between steps. For PC-PC interactions, Particle mesh Ewald (PME) was used with grid-size spacing of 1 Å, a relative tolerance of 10, and a 12-Å cutoff with 10 Å switching for the Lennard-Jones (LJ) interactions. For higher MTP interactions, power-law dependent switching was employed. Protein stability differences ($\Delta\Delta G_{stab}$) were calculated according to $\Delta G_2 - \Delta G_1$, as shown in the thermodynamic cycle in Figure S1a, where $\Delta G_1$ and $\Delta G_2$ are the free energies of mutating the benzene ring $C_6H_6$ into $C_6H_5X$ (X is the halogenic substitution), in aqueous phase, either as a single residue or in the protein, respectively. Destabilizing mutations would thus have a positive $\Delta\Delta G$. For the experimental values (below) a sign convention was adopted such that a positive $\Delta\Delta G$ indicates that the mutant is less stable than the WT protein. Each calculation of $\Delta\Delta G_{stab}$ entailed a *forward* (WT → mutant) and a *backward* (mutant → WT) simulations, which provided lower and upper bounds to the free-energy difference, respectively. These simulations were repeated 5 times each. The results reported in Table 1 (main text) represent the mean of 5 trials.

**Quantum-Chemical Calculations.** The electrostatic surface potentials (incorporated electron density) were obtained from *ab initio* calculations and mapped at the $10^{-3}$ $ea^{-3}$ isodensity surface using Gaussview5. All *ab initio* calculations were carried out with Gaussian09[10] at the MP2 level with the aug-cc-pVDZ basis set, and optimized structures were used.

**Synthesis of Insulin Analogs.** Insulin analogs were prepared by trypsin-catalyzed semi-synthesis.[15] This protocol employs (i) a synthetic octapeptide representing residues (N)- GF*FYT<u>KP</u>T (including modified residue (F*) and "KP" substitutions (underlined) and (ii) truncated analog *des*-octapeptide[B23-B30]-insulin. Because the octapeptide differs from the wild-type B23-B30 sequence (GFFYT*PK*T) by interchange of $Pro^{B28}$ and $Lys^{B29}$ (italics), protection of the lysine ε-amino group is not required during trypsin treatment. In brief, des-octapeptide (15 mg) and octapeptide (15 mg) were dissolved in a mixture of dimethylacetamide/1,4-butandiol/0.2 M Tris acetate (pH 8) containing 10 mM calcium acetate and 1 mM ethylene diamine tetra-acetic acid (EDTA) (35:35:30, *v/v*, 0.4 mL). The final pH was adjusted to 7.0 with 10 µL of N-methylmorpholine. The solution was cooled to 12 ºC, and 1.5 mg of TPCK-trypsin was added and incubated for 2 days at 12 ºC. An additional 1.5 mg of trypsin was added after 24 hr. The reaction was acidified with 0.1% trifluoroacetic acid (TFA) and purified by preparative reverse-phase HPLC (C4). Mass spectrometry using matrix-assisted laser desorption/ionization time-of-flight (MALDI-TOF-TOF; Applied Biosystems, Foster City, CA) in each case gave expected values. The protocol for solid-phase synthesis is as described.[16] (Fmoc)-protected Phe analogs were purchased from Chem-Impex International (Wood Dale, IL).

**CD Spectroscopy.** Spectra were obtained at 25 ºC using an Aviv spectropolarimeter.[17] Samples contained *ca*. 25 µM insulin analog in 50 mM potassium phosphate (pH 7.4); samples were diluted to 5 µM for guanidine-induced denaturation studies at 25 ºC. To extract free energies of unfolding ($\Delta G_u$), denaturation transitions were fitted by non-linear least squares to a two-state model as described.[18]

**Receptor-Binding Assays.** Relative activity is defined as the ratio of hormone-receptor association constants relative to human insulin. Association constants were determined by a competitive-displacement assay using $^{125}$I-$Tyr^{A14}$-insulin (kindly provided by Novo-Nordisk) and solubilized insulin receptor (isoform B) in a microtiter plate antibody capture assay as described[19] transfected receptors were tagged at the C-terminus by a triple repeat of the FLAG epitope (DYKDDDDK). Microtiter plates were coated by an anti-FLAG M2 monoclonal antibody (Sigma). The percentage of tracer bound in the absence of competing ligand was <15% to avoid ligand-depletion artefacts. Binding data were analyzed by nonlinear regression using a heterologous competition model.[20]

**Cell Culture.** Signaling activities of the analogs were tested in L6 (rat myoblast cell line) with exogenous human IR expression.[21] L6-IRA cells were cultured in DMEM medium supplemented with 10% FBS, and G418 providing selection condition. 24-hours serum-starving protocol using culture medium except FBS was applied after reaching 70-75% confluence (approximately $0.8X10^6$ cells per well). After the starving, serum-free medium containing 50 nM of tested insulin analogs were added in each well simultaneously. Time of treatment was 15 mins and insulin-containing medium was removed, followed by the cell-lysis protocol using RIPA buffer with protease and phosphatase inhibitor (Roche). Cell lysates were collected and BCA assay was used to measure protein concentrations for immunoblotting study.

**Western Blot**. Samples for the Western-blot contained approximately 5 µg of protein in each loading well. Lysates were dissolved in Laemmli buffer (Bio-Rad) with 10% beta-mercaptoethanol and heated at 50 °C for 10 minutes, and then centrifuged at 10,000 RPM for 1 minute. Samples were loaded into 10% Mini-PROTEAN TGX gels (Bio-Rad).

Proteins were transferred to PVDF membrane, and then blocked in 5% BSA for 1 hour. Membranes were incubated overnight at 4 °C cold room with Insulin Receptor β (4B8) Rabbit mAb (Cell Signal) or an equal mixture of Phospho-Insulin Receptor β (Tyr1150/1151) Rabbit mAb (Cell Signal); Phospho-Insulin Receptor (Tyr1158) Polyclonal Antibody (Thermo Fisher Scientific); Phospho-Insulin Receptor (Tyr1334) Polyclonal Antibody (Thermo Fisher Scientific); Phospho-Insulin Receptor β (Tyr1345)

Rabbit mAb (Cell Signal); and Anti-Insulin Receptor (phospho Y972) antibody (Abcam). Dilutions for these antibodies were 1:5000 in 5% BSA. After the primary antibody incubation, membranes were washed with TBS with 0.1% Tween-20 and incubated in Goat Anti-rabbit HRP conjugated secondary antibody diluted 1:10,000 in 5% for 1-2 hours at room temperature. Membranes were washed three times with TBS with 0.1% Tween-20 and incubated in HRP Substrate (EMD Millipore) for 45s and developed.

**Realtime-qPCR.** L6-IRA cells treated for 8 h were collected, and the transcriptional activation of cyclin D1 and the transcriptional repression of cyclin G2 were assessed by rt-qPCR. Sample preparation were followed the instruction of One-Step rt-PCR reagent kits as described by the vendor (Bio-rad).

The following sets of primers were used.

Cyclin D1: 5′-GCCGAGTGGAAACTTTTGTCG-3’ and 5′-CGGGAAGCGTGTACTTATCCT-3’
Cyclin G2: 5’- GCAAGAAAAGAAGCCAAGCT-3’ and 5’-
TGACCAAGAGGCAAAATAAAATCAA-3’
GAPDH: 5’-GACATGCCGCCTGGAGAA-3’ and 5’- GCCCAGGATGCCCTTTAGT-3’
TFIID: 5’-CTGAGGGGGCAATGTCTAAC-3’ and 5’-GGGCAGCTAGTGAGATGAGC-3’.

**Rat studies.** Groups of male Lewis rats rendered diabetic by pancreatic β-cell toxin streptozotocin were utilized. Analyses were conducted following intravenous (i.v.) bolus injection. Rats were injected at $t$ = 0 and then every 10 min for the 1st hour, every 20 min for the 2nd hour, every 30 min during the 3rd hour and then once per hour for the remainder of the experiment. The doses of insulin analogs administered were 1.7 nmol per 300-gram rat (n=6) and 2.6 nmol per 300-gram rat (n=5). Measurements of [glucose] were made using a clinical glucometer (Hypoguard Advance Micro-Draw meter).

##### Assessment of fibril formation

**Accelerated protocol.** The physical stability of the analogs was assessed by their propensity to form fibrils. To estimate lag times prior to onset of fibril formation, halogen containing KP-insulins or control KP-insulin were made 60 μM in 1x phosphate-buffered saline (pH 7.4) containing 0.02 % sodium azide as an antimicrobial agent in presence of 16 μM thioflavin T (ThT). The insulin solutions in replicates (n = 8) were plated in Costar® plate and continuously shaked at 37 °C and at 960 rpm in a BioTek plate reader. The time of initial THT fluorescence was recorded as fibrillation lag time.

**Gentle rocking protocol.** The insulin solutions at 60 μM concentration in phosphate-buffered saline (pH 7.4) containing 0.1% sodium azide with gentle rocking at 37 °C were incubated in glass vials containing a liquid/ air interface. Aliquots were taken at regular intervals and frozen to enable analysis of thioflavin T fluorescence at the end of the assay, terminated on visual appearance of cloudiness in the solution.

#### II. Supplemental Simulations

***Flexibility of B-chain C-terminal segment (B24-B29) depends on specific B24 modification.*** To relate predicted changes in thermodynamic stability to potential conformational changes, ten additional MD simulations were carried out: the WT monomer; F-Phe$^{B24}$ in positions 2, 3, 4, 5 and 6; and the Cl-Phe$^{B24}$ and Br- Phe$^{B24}$ variants in the two *ortho* positions (2 and 6). One independent trajectory (10 ns) was run for each system. The analysis focused on displacement of the C-terminal segment of the B chain due to its immediate relevance to the mechanism of receptor binding.[22]

C-terminal B-chain opening was first investigated for the fluorinated Phe$^{B24}$ monomers. Figure S2 shows the role of fluorination on the conformational dynamics of the B-chain C-terminal segment. The evolution of the distance between the N-backbone atom of residue A1 and the C=O backbone atom of residue B30 was monitored ("$d_{A1\text{-}B30}$"; Figure S2a). Figure S2b shows RMSD values relative to the starting X-ray conformation (residues B18-B30). Figure S2c illustrates the chain movement by superposition of the reference PDB structure (4INS; in red) to 20 frames as sampled along the MD simulation. This comparison provides an overview of the displacement of the B-chain residues with respect to Gly$^{A1}$ (Figure S2a). For the WT monomer the distance progressively increased from around 10 Å to 15 Å during the MD simulation. Each of the fluorinated proteins exhibited larger distance fluctuations ranging from 10 to 22 Å. Nevertheless, in some cases (4F-Phe$^{B24}$, 5F-Phe$^{B24}$ and especially for 2F-Phe$^{B24}$) the distance also decreased to 2.5-3 Å in the course of the trajectories. For 2F-Phe$^{B24}$ variant (lower panel in Figure S2a, black line) $d_{A1\text{-}B30}$ decreased from *ca.* 12.5 to 2.5 Å in the first 3 ns and then maintained this closed conformation for the majority of the simulation.

$\alpha$-Helical conformations were stable in each MD simulation. The RMSD from the starting conformation is *ca.* 1 Å (data not shown). In contrast, the N-terminal and the C-terminal ends of B-chain were flexible. For WT insulin, RMSD values of the B-chain C-terminal segment increased from *ca.* 1.0 to 1.5 Å during the first 2 ns of the MD simulation and then stabilized around 2.5 Å. The flouro-Phe$^{B24}$ analogs each exhibited higher RMSD values, beginning at 2 Å and increasing to 4 Å (after 1 ns for F in *para*, in *meta* and in *ortho*-6 positions or after 4 ns for F in *ortho*-2; Figure S2b).

Superposition of frames sampled from the 10 ns MD trajectories (shown in Figure S2c in relation to the reference PDB monomer 4INS; red) enabled monitoring of conformational changes of the B-chain C-terminal segment. These structures reflect the above trends in $d_{A1\text{-}B30}$ values (Figure S2a), including the "locked" conformation of the 2-F-Phe$^{B24}$ analog. Profound differences were seen in the simulations of corresponding 2-Cl-Phe$^{B24}$ and 2-Br-Phe$^{B24}$ analogs (Figure S3). Whereas 2F led to occlusion of the A1-A3 region, this receptor-binding surface was in part re-opened by 2-Cl-Phe$^{B24}$; 2-Br-Phe$^{B24}$ was associated with an entirely new conformation. Similar profound differences were also observed for *ortho*-6 halogenated analogs (Figure S4). These effects are discussed below in relation to backbone dihedral-angle distributions.

***B24 modifications introduce new local interactions.*** The above MD simulations suggested that the B-chain C-terminal segment can differ in flexibility depending on the modification. To explore the origins of these effects, the backbone dihedral angles of X-Phe$^{B24}$, Phe$^{B25}$ and Tyr$^{B26}$ were considered next.

*Phi* and *psi* dihedral-angle distributions (respectively designated $P(\Phi)$ and $P(\Psi)$) for X-Phe$^{B24}$ variants are shown in Figures S6-S7 and Figure 4a (main text) relative to WT. Figure S6 presents $P(\Phi)$ and $P(\Psi)$ of F- Phe$^{B24}$, Phe$^{B25}$ and Tyr$^{B26}$ for the five fluorinated analogs. Whereas in each case B24 $P(\Psi)$, B26 $P(\Psi)$ and B24 $P(\Phi)$ were similar to WT, marked differences were observed in B26 $P(\Phi)$ and in both B25 $P(\Psi)$ $P(\Phi)$ specifically for 2- and 6-F.

(i) **2F-Phe$^{B24}$ simulation**. The B-chain bent away from its WT orientation at Tyr$^{B26}$ (Figure S6, top panel, solid red line); this was apparently due to an intra-residue interaction between the F and the B24 backbone N-H (Figure 3a in main text). This interaction required a change in B24 Φ angle from -150° to -125°, in turn associated with a change in B25 Φ from -100°

to -160° (middle panel) and leading to two B26 conformational states, *open* and *closed* (the former with Φ = -150° and latter at Φ = -50°; *versus* -125° for WT). The acute Φ angle (-50°) oriented the C-terminal B-chain segment (B26-B30) toward the A chain. In this closed conformation $Thr^{B30}$ engages both the α-amino group of $Gly^{A1}$ and the δ–$NH_2$ of $Glu^{A17}$ (Figure 3b in main text).

(ii) **6F-$Phe^{B24}$ simulation**. Although B24 main-chain dihedral angles were not affected (Figure S6, bottom panel, dashed red line), B25 Φ decreased by 50° relative to WT (middle panel), and a new *P*(Ψ) appeared at 10° with the same probability as 125°. This was associated with a shift in B26 *P*(Ψ)from -125° to -60° (upper panel). This concerted set of conformational changes presumably reflected electrostatic interactions between the 6-F and the amide–NH main-chain atoms of $Phe^{B25}$ and $Tyr^{B26}$ (Figure S5).

Figure 4a of the main text focuses on the three modifications at position 2 (X=F, Cl, Br). Comparison of the *P*(Φ) and *P*(Ψ) distributions of $Phe^{B24}$, $Phe^{B25}$ and $Tyr^{B26}$ highlighted local changes in the orientation and interactions of nearby side chains.

(iii) **2Cl-$Phe^{B24}$ simulation**. Like 2-F, 2-Br was associated with a shift the *P*(Φ) peak from -150° (WT) to -125°, whereas 2-Cl was associated with a wider distribution ranging from -125° to -80°. Distinct patterns of transmitted conformational changes were observed. Whereas 2-Cl spared the B25 *P*(Φ) distribution, 2-F and 2-Br were associated with respective shifts in B25 *P*(Φ) peak to 150°and to -60°. 2Cl-, the emergence of a new *P*(Ψ), compared to the WT (≅ 125°), ranging from -50° to 0°.

(iv) **2Br-$Phe^{B24}$ simulation**. For $Tyr^{B26}$, 2Cl- invokes similar modification to *P*(Φ) as 2F-, but with different probabilities, while 2Br- leads to the appearance of a wide distribution ranging from -100° to -60°. Br also induces a shift of the $Tyr^{B26}$ *P*(Ψ) peak from ≅ 140° (for WT, 2F- and 2Cl-) to -40°.

The dynamic behavior of the B-chain C-terminal segment in the 2Cl-$Phe^{B24}$ mutant can be related to the Cl atom interacting with positive and negative sites of the protein via both its electron rich $\delta^-$ and electron deficient $\delta^+$ regions (Figure 3c in main text). The $\delta^+$ region forms a σ-hole bond with the carbonyl–O backbone of $Gly^{B20}$ and the $\delta^-$ region interacts with the amide–NH backbone atoms of $Gly^{B23}$ and $Phe^{B24}$, between which it alternates. Due to the Cl--O σ-hole bond, the $Phe^{B24}$ residue is pulled towards the U-turn which explains the B-chain bending away from A-chain (Figure 3d in main text). For the 2Br-$Phe^{B24}$ mutant, the large vdW radius of Br allows the formation of an H-bond with the amide–NH backbone atom of $Phe^{B24}$ and $Tyr^{B16}$, respectively (Figure 3e in main text). Br bridges between the β-strand and the α-helix of chain-B bringing them closer to each other at the $Tyr^{B16}$ – $Phe^{B24}$ level, which causes the U-turn to tighten. The backbone carbonyl-O of $Gly^{B20}$ is then pushed to larger distances from Br and no σ-hole bond is formed.

In Figure S7 is shown a comparison of $Phe^{B24}$ halogenation in the *ortho*-6 position relative to WT; the *P*(Φ) peak of $Phe^{B24}$ shifts from -150° to -135° and -115° when passing to Cl and Br, respectively. In this position, the halogens are exposed to the backbone carbonyl-O of $Phe^{B24}$ and thus the decrease of *P*(Φ) when passing from F to Cl to Br is dependent on the size of the halogen introduced. The *P*(Ψ) of $Phe^{B24}$ shows in the case of Br the emergence of a new state at 125°. For $Phe^{B25}$, 6F- and 6Cl- shifts the *P*(Φ) peak from -100° to -135° and -140° respectively, while 6Br- broadens the angular distribution peak from -100° to -75°, and shifts the *P*(Ψ) from 75° to 15° and -45°. The effect of the 6X- halogenations on $Tyr^{B26}$ affects only *P*(Φ) , where F, Cl and Br shift the peak from -125° to -60° but with a lower probability in the case of Br. The 6Br- mutation also shows the emergence of a second peak at -145° with a higher probability than the peak at -60°. The dihedral angular variations induced by the 6Cl- mutation can be explained by the halogen bond formation between the latter and the amide–NH backbone atoms of $Phe^{B25}$ and $Tyr^{B26}$, as in 6F- (Figure S5a,b). However, the accommodation of the larger vdW radius of Cl, compared to F, induces the emergence of new *P*(Φ) and *P*(Ψ) for $Phe^{B25}$. The largest change in *P*(Φ) and *P*(Ψ) emerge from the 6Br- halogenation. In this case, there is a σ-hole bond formation between the $\delta^+$

region of Br and the carbonyl-O backbone of $Phe^{B24}$ in a first step (Figure S5c), and in a second step the B24 ring flips and Br interacts with the amide–NH backbone atom via its $\delta^{-}$ region (Figure S5d); this observation explains the emergence of two different peaks in the dihedral angle distributions.

***Enthalpy-Entropy Compensation Explains Stabilization of 2X versus 6X.*** Interaction energies ($\Delta E_{int}$) were analyzed for halogen substituents at positions 2 and 6 (Figure S8 and S9, respectively), as calculated from the above 10-ns MD simulations. The analyses focused on three sets of interactions:

- between the B24 side chain and the A-chain, and
- between the B24 side chain and the B-chain.

Non-bonded energy contributions were decomposed into an electrostatic ($\Delta E_{elec}$; red lines in the figures) and a van der Waals ($\Delta E_{vdW}$; green lines) component.

**Position 2.** Relative to WT, halogen atoms at position 2 generally increased interaction energies between the modified side chain and the A-chain (Figure S8a).

(i) **2-F-Phe$^{B24}$.** $\Delta E_{int}$ increased from -2 (WT) to *ca.* -4 kcal/mol (Figure S8a); the interaction energy with the B-chain decreased by ~ 5 kcal/mol (Figure S8b). In each case the total energy was mirrored by its electrostatic contribution $\Delta E_{elec}$ (red lines). Thus, 2-F increased (decreased) the interaction energy with A-chain (B-chain) residues. This trend was associated with the movement of the B-chain C-terminus towards the N-terminal A-chain $\alpha$-helix and away from the central B-chain $\alpha$-helix (Figure S3c).

(ii) **2-Cl-Phe$^{B24}$.** The increased $\Delta E_{int}$ was driven almost entirely by increased interactions with the B-chain (~ 5 kcal/mol; Figure S8b) whereas $\Delta E_{int}$ with the A-chain was not significant (Figure S8a) in accordance with structural trends opposite to those induced by 2-F (Figure S3c). A striking implication is that 2-F and 2-Br can each achieve net enthalpic stabilization via distinct atomic mechanisms.

(iii) **2-Br-Phe$^{B24}$.** The pattern of interaction energies was similar to 2-F. The larger structural excursions in this case (Figure S3c) were associated with larger fluctuations in interaction energies. 2-Br enhanced $\Delta E_{int}$ between B24 and A-chain residues after 1 ns of MD simulation (ranging between -2 and -8 kcal/mol; Figure S8a) and $\Delta E_{int}$ between B24 and B-chain residues (ranging from -7.5 to 7.5 kcal/mol; Figure S8b). This trend was associated with distinct structural changes in the C-terminal B-chain segment, arising from $\Phi$ and $\Psi$ angular variations of the base residues $Phe^{B24}$, $Phe^{B25}$ and $Tyr^{B26}$ (Figure 4a in main text). These changes enabled the C-terminal B-chain residues to engage with both the N-terminal A-chain $\alpha$-helix and central B-chain $\alpha$-helix (Figure S3c).

Relative to 2-F, the total energy profiles in the larger halogens were primarily driven by its $\Delta E_{vdW}$ contribution. The more substantial fluctuations of $\Delta E_{int}$ in the 2-Cl and 2-Br simulations were each associated with greater flexibility of the B-chain C-terminal segment (Figure S3).

**Position 6.** Electrostatic terms made a leading contribution at position 6; $\Delta E_{vdW}$ terms were similar among the halogens (Figure S9b). The halogen substituents at position 6 induced large energy fluctuations associated with flexibility of the C-terminal B-chain segment (Figure S4). This trend was more pronounced than in the 2-X simulations as observed by RMSD values and $d_{A1\text{-}B30}$ evaluations (Figure S4a,b)). These findings predict increased changes in configurational entropy relative to the 2-X simulations.

(i) **6-F-Phe$^{B24}$.** The C-terminal B-chain segment alternated between two states, one leaning toward the A-chain and the other leaning away (Figure S4c). These two states underlay the increased $\Delta E_{int}$ between 6F-$Phe^{B24}$ and residues in both chains (Figure S9a,b).

(ii) **6-Cl-Phe$^{B24}$.** $\Delta E_{int}$ was increased between 6-Cl-$Phe^{B24}$ and the-A chain but decreased between $Phe^{B24}$ and the B-chain. This trend was associated with preferred orientation of the C-terminal B-chain segment toward the A-chain.

(iii) **6-Br-Phe$^{B24}$.** Changes in $\Delta E_{int}$ terms and structural displacements of the B-chain C-terminal segment were opposite to that of 6-Cl (Figure S9a,b and Figure S4c).

Residues in insulin affected by the B24 modifications at positions 2 or 6 were identified (Tables S1 and S2, respectively). These tables provide a comparison of vdW ($\Delta E_{vdW}$) and electrostatic ($\Delta E_{elec}$) energy fluctuation differences of backbone (bb) and side-chain (sc) atoms relative to WT, as calculated from the 10-ns MD simulation. In addition to the modified residue itself (B24), major enthalpic fluctuations were found to involve residues Gly$^{A1}$, Glu$^{A4}$, Glu$^{A17}$, Tyr$^{A19}$, Cys$^{A20}$, Gln$^{A21}$, Val$^{B12}$, Leu$^{B15}$, Tyr$^{B16}$, Cys$^{B19}$, Gly$^{B20}$, Arg$^{B22}$, Gly$^{B23}$, Phe$^{B25}$, and Tyr$^{B26}$. Most of these residues are not in direct contact with Phe$^{B24}$.

These results are in accordance with the observed interactions (Figure 3 in main text and Figure S5) and with the above analysis of segmental conformational changes (Figure S2-S7 and Figures 4a). Favorable changes in enthalpy were dipole dependent in the series (F > Cl > Br). Predicted effects are more marked at position 2 (-9.58 > -8.94> -8.81 kcal/mole) than at position 6 (-7.08 > -3.15 > -2.67 kcal/mole). As indicated by the free-energy calculations in the main text (Table 1), changes in entropy following modification at position 2 cannot obviate this highly increased enthalpy, whereas at position 6 the enthalpic terms are substantially offset by compensating entropic terms. These simulations thus predict that *ortho*-X-Phe$^{B24}$ insulin analogs will predominantly occupy the 2 position in a conformational equilibrium related by $\chi_2$ angle.

***B24 modifications affect interactions within the hormone-receptor complex.*** Inspection of recent structures of insulin- μIR[22] and insulin-ectodomain complexes[23] revealed packing defects adjoining B24 ring positions 2 and 6 (wider at 2), whereas ring positions 3-5 are tightly packed. For this reason (and because of the enhanced stabilities of the free hormone variants), the present MD simulations of variant μIR complexes focused on *ortho* substitutions; free-energy calculations were not performed.

Analysis of the interaction energies (enthalpies) between the hormone and μIR were undertaken based on 1-ns MD simulations of the WT complex and the following six halogen derivatives: 2-F, 2-Cl, 2-Br, 6-F, 6-Cl, and 6-Br. In addition, 2-$CH_3$- and 6-$CH_3$ Phe$^{B24}$ analogs were simulated as non-halogen controls. The energy decomposition analysis of these interaction energies (extracted from the MD simulations) demonstrated the greater general importance of vdW contributions in the pocket relative to electrostatic terms. The larger substituents in general mitigated the packing defects adjoining ring positions 2 or 6. However it was more favorable at ring position 2 than 6 (since the packing defect adjoining B24 ring position 2 is wider than position 6), as can be seen by the higher vdW energy distribution for Cl, Br and $CH_3$ in position 2 (Figure 10a) compared to position 6 (Figure 10b). The electrostatic properties of the halogenic substituents are generally unfavourable within the native B24-binding pocket. Indeed, the larger the dipole (in order F>Cl>Br>$CH_3$=0), the less favorable is the electrostatic component of binding (Figure S10). The observed trend in experimental affinities (F<Cl~Br<$CH_3$; Table S3) is in accordance with these calculations. However, while the enthalpy contribution suggested that certain halogen derivatives may bind more tightly than WT (2-Cl, 2-Br, 6-Br, and 6-Cl), this was not observed experimentally. We speculate that the resulting entropy change was higher than the enthalpy gain due to the frustration induced by the presence of the dipole in the pocket. A signature of this frustration was provided by the height and width of the probability distributions (Figure S10). This signature was not seen in the WT simulation nor in the control (2/6)-$CH_3$-Phe$^{B24}$ simulation (See Figure S10).

a)
b)

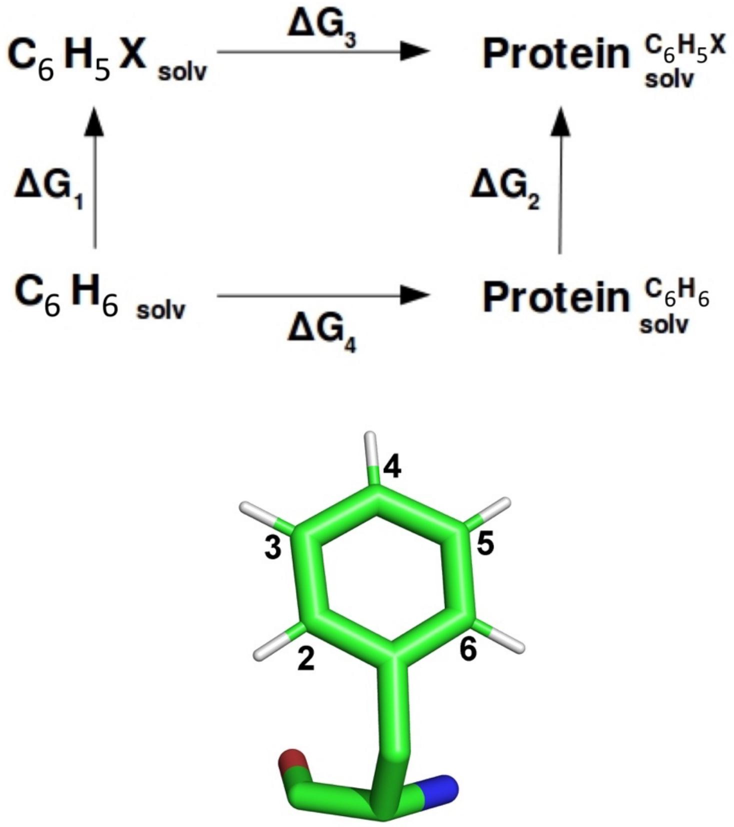


**Figure S1.** a) Thermodynamic cycle used to calculate the free energy differences in stability ($\Delta\Delta G_{\text{stab}}$) between WT and B24-modified proteins: X designates substitution of an aromatic hydrogen by a halogen atom. b) Phe side chain with carbon-atom positions 2-6 labeled (*ortho*-[2/6], *meta*-[3/5], *para*-4).

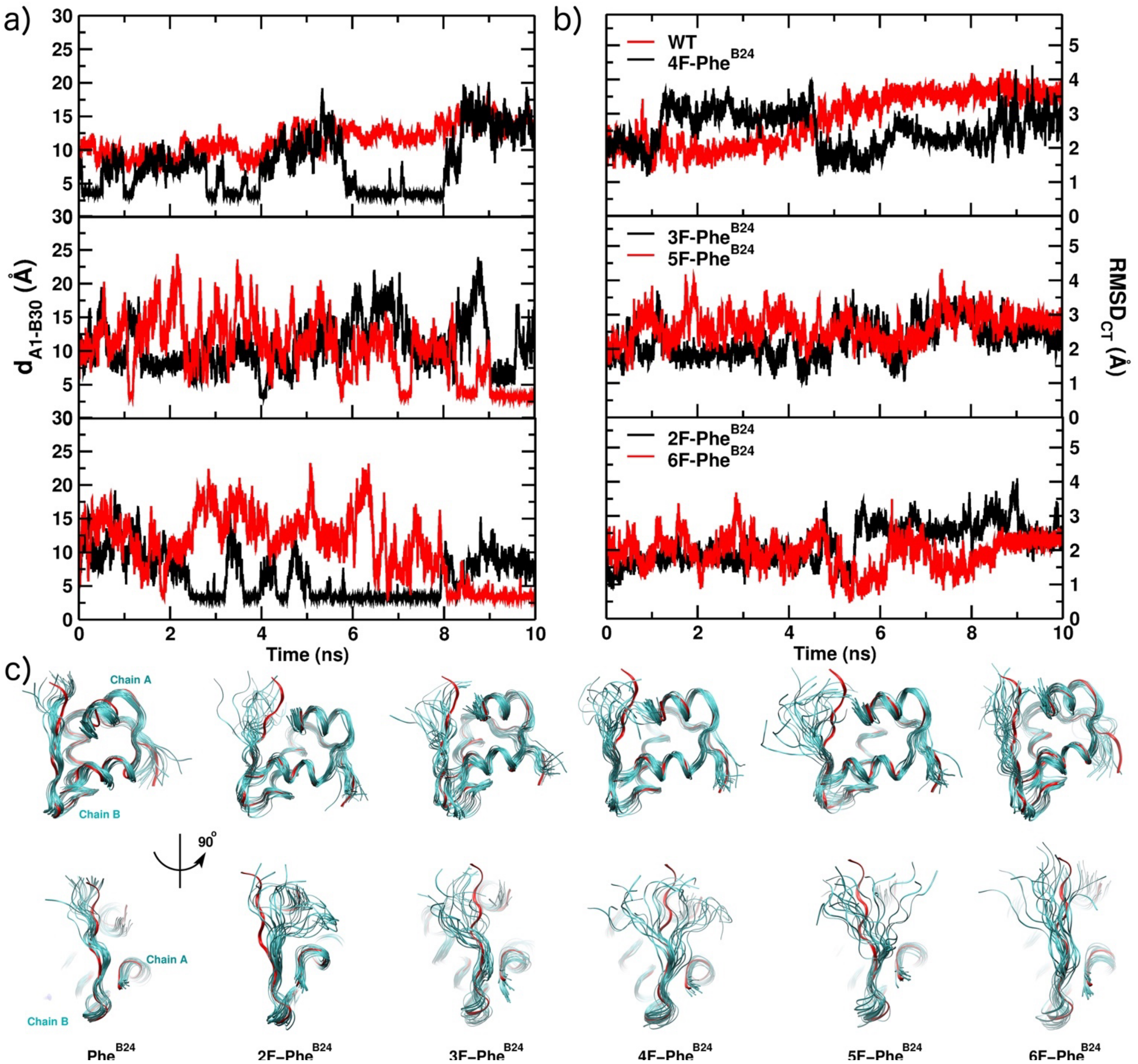


**Figure S2.** B24 fluorination-induced conformational changes of the B-chain C terminal segment. a) Evolution of the distance between the N backbone atom of residue A1 and the C backbone atom of residue B30 during 10 ns of MD simulation; b) RMSD from the starting conformation of the insulin WT monomer during 10 ns of the MD simulation, calculated for the C-terminal B-chain residues (B18-B30); c) Overlay of 20 structures sampled from the 10-ns MD simulation trajectories. The frames are separated by 0.5 ns, and the reference X-ray structure of the INS monomer is shown in red. In the bottom row the B chain is facing the viewer.

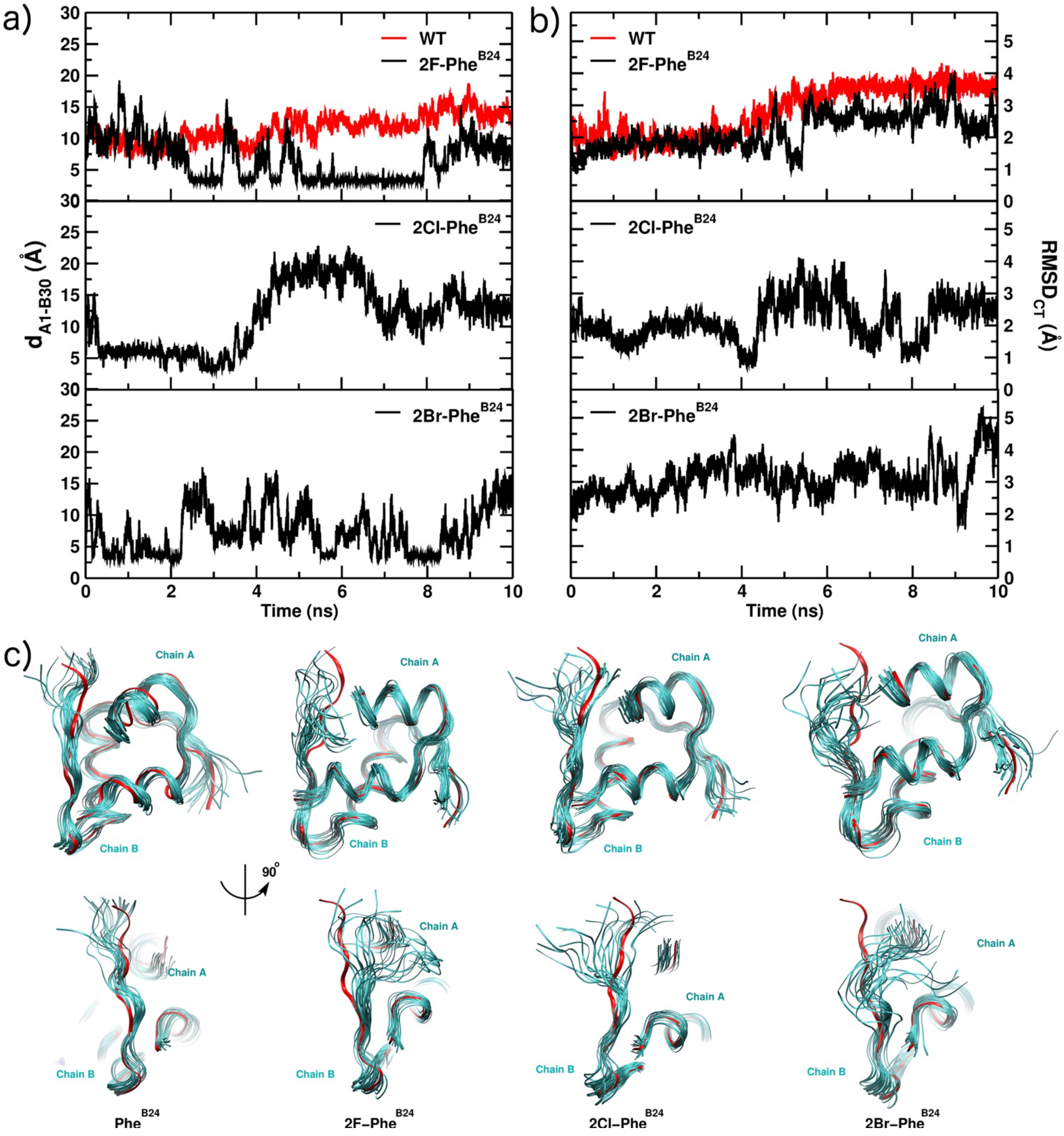


**Figure S3.** Monitoring the "2X-B24"-induced conformational changes of the B-chain C-terminal segment. 2X represents F, Cl and Br at the *ortho*-2 position. a) Evolution of the distance between the N backbone atom of residue A1 and the C backbone atom of residue B30 during 10 ns of MD simulation; b) RMSD from the starting conformation of the WT insulin monomer during 10 ns of the MD simulation, calculated for the C-terminal B-chain residues (B18-B30); c) Overlay of 20 structures sampled from the 10-ns MD simulation trajectories. The frames are separated by 0.5 ns, and the reference X-ray structure of the INS monomer is shown in red. In the bottom row the B chain is facing the viewer.

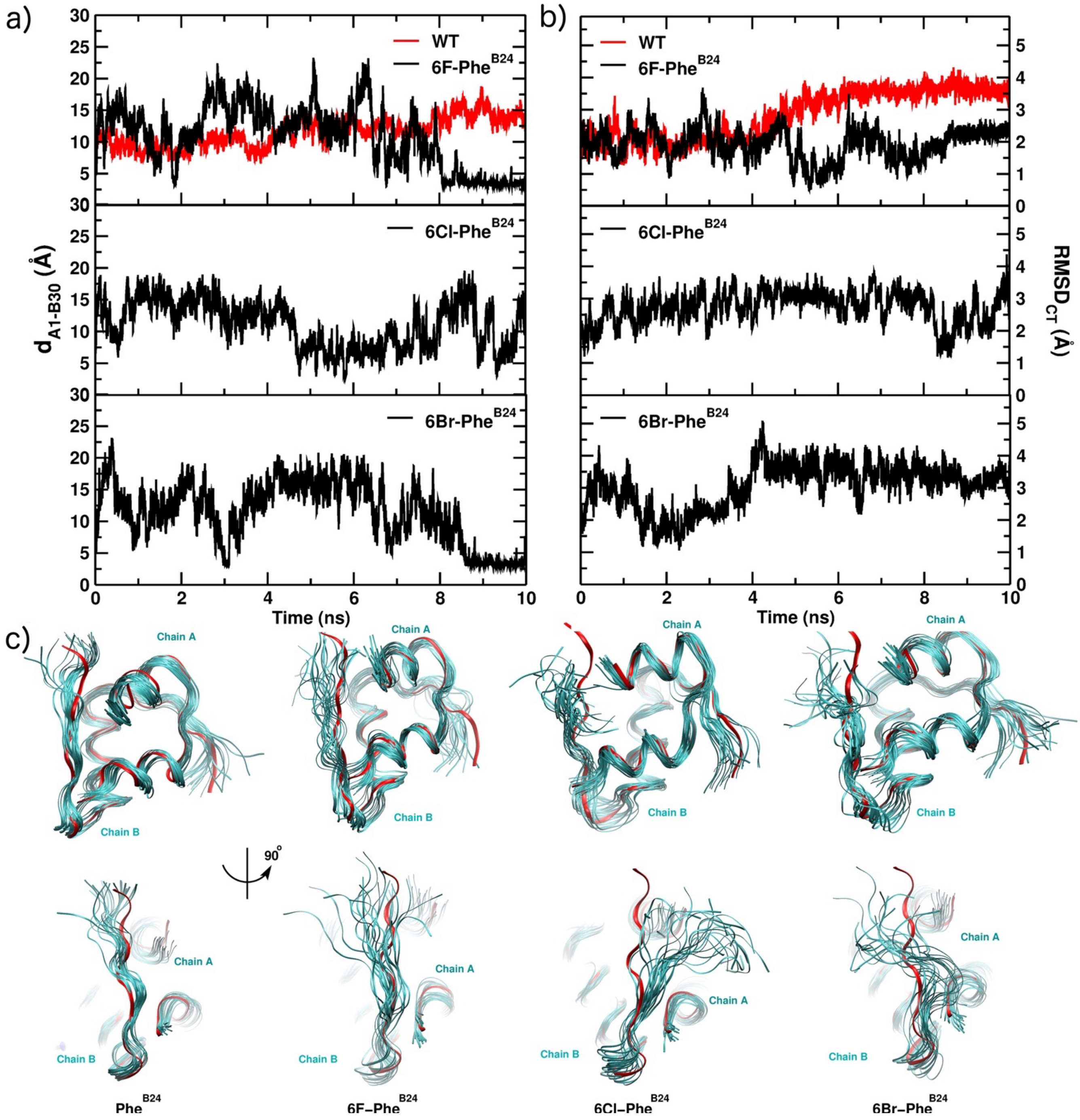


**Figure S4.** Monitoring the "6X-B24"-induced conformational changes of the B-chain C terminus. 6X represents F, Cl and Br at the *ortho*-6 position. a) Evolution of the distance between the N backbone atom of residue A1 and the C backbone atom of residue B30 during 10 ns of MD simulation; b) RMSD from the starting conformation of the WT insulin monomer during 10 ns of the MD simulation, calculated for the C-terminal B-chain residues (B18-B30); c) Overlay of 20 structures sampled from the 10 ns MD simulation trajectories. The frames are separated by 0.5 ns, and the reference X-ray structure of the INS monomer is in red. In the bottom row the B chain is facing the viewer.

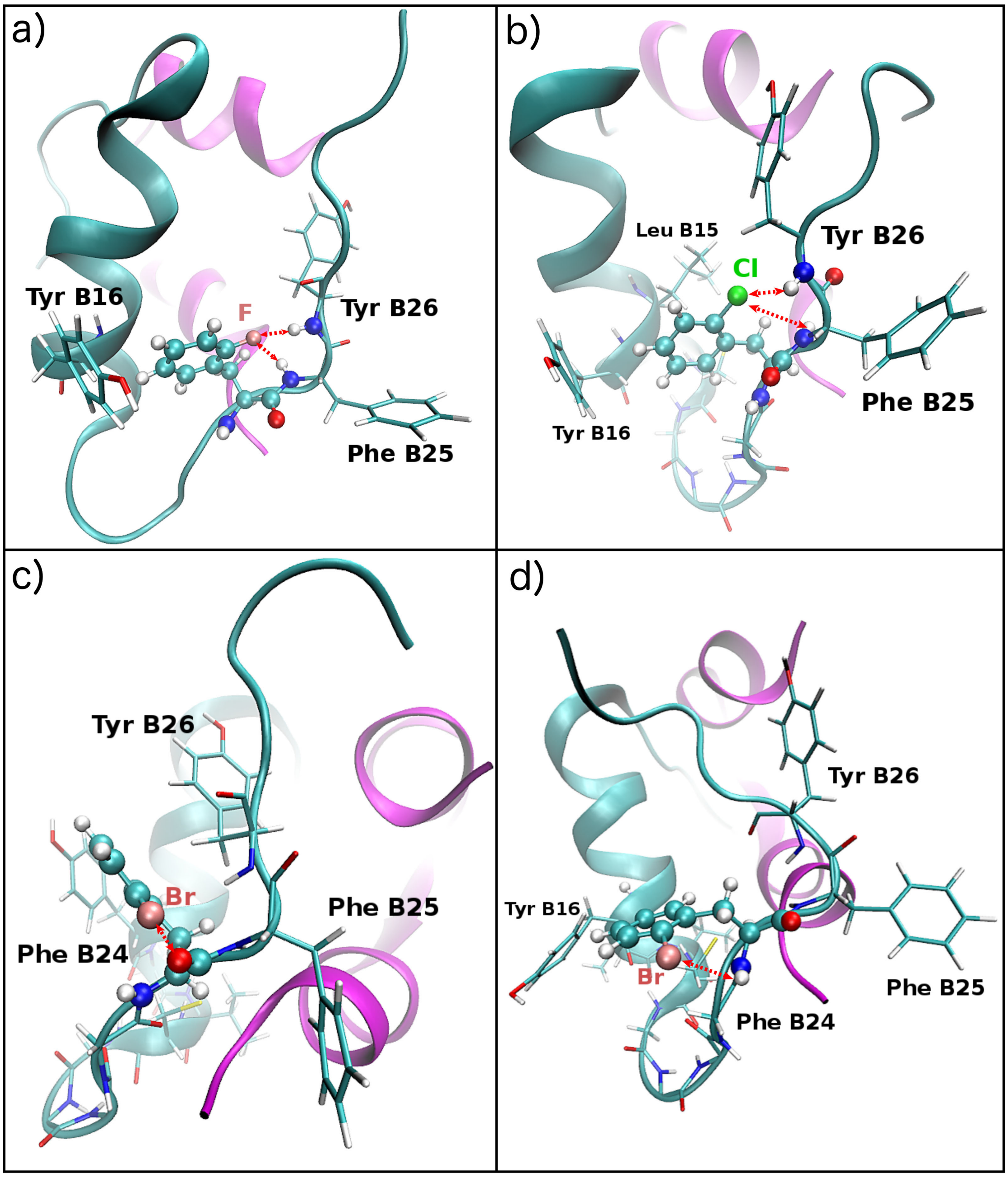


**Figure S5.** Structures of halogenated 6-X-Phe$^{B24}$ insulin monomers. a) T-shaped structure of the "6-X-Phe$^{B24}$" – Tyr$^{B16}$ $\pi$-stacking; interactions of fluorine with both –NH backbone of Phe$^{B25}$ and Tyr$^{B26}$. b) Interactions of Cl with the –NH backbone atoms of Phe$^{B25}$ and Tyr$^{B26}$, and repulsion of the Tyr$^{B26}$ backbone O. c) The figure illustrates the formation of a sigma-hole interaction between the $\delta^{+}$ region of Br and the O backbone atom of 6-Br-Phe$^{B24}$. d) B24 ring flips and Br interacts with the –NH backbone atom. Red dashed arrows indicate the hydrogen/halogen-bonds.

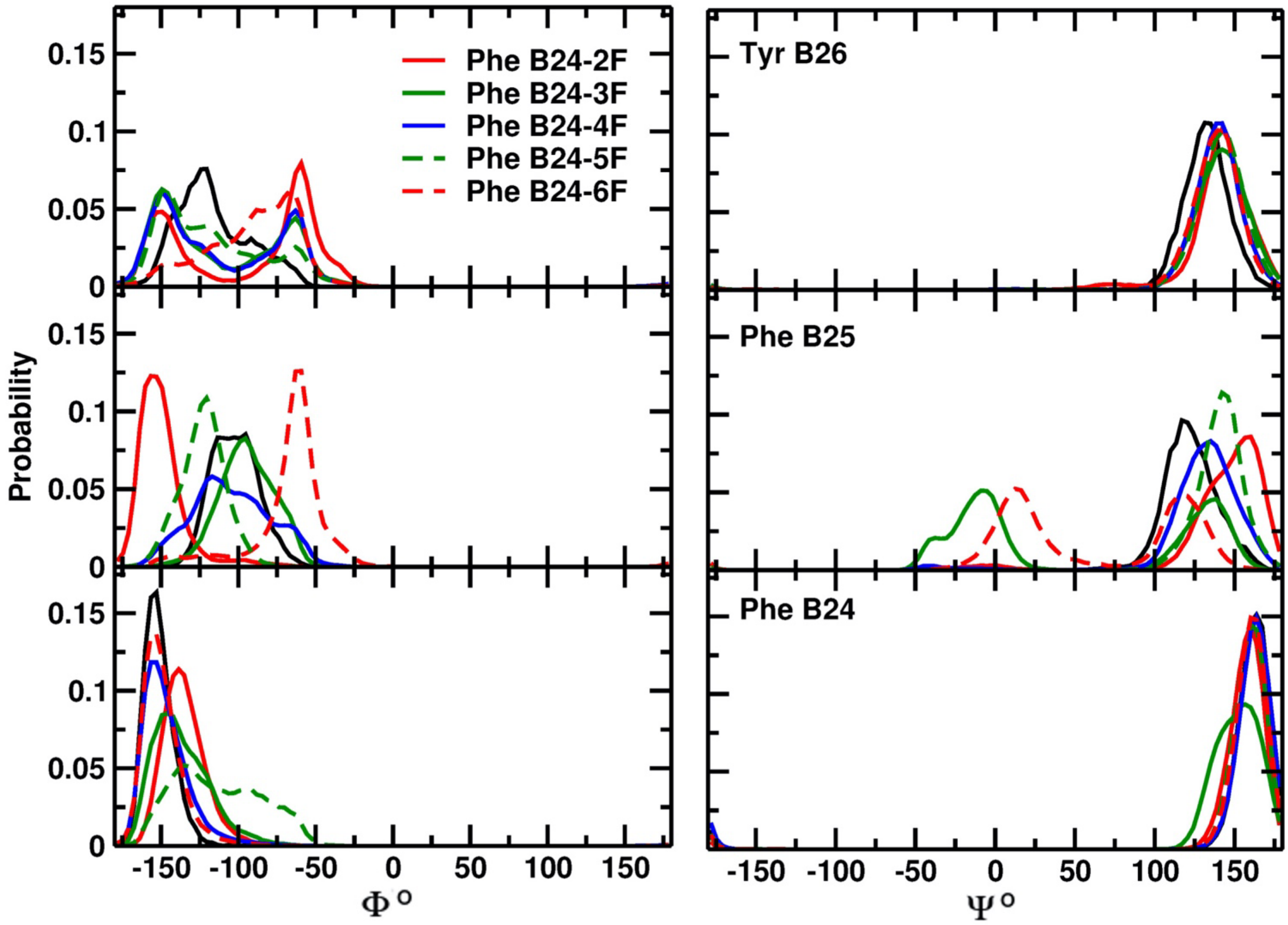


**Figure S6.** Dihedral angles distribution of Phe[B24], Phe[B25] and Tyr[B26] for each of the five possible Phe[B24] fluorination sites in the aromatic ring.

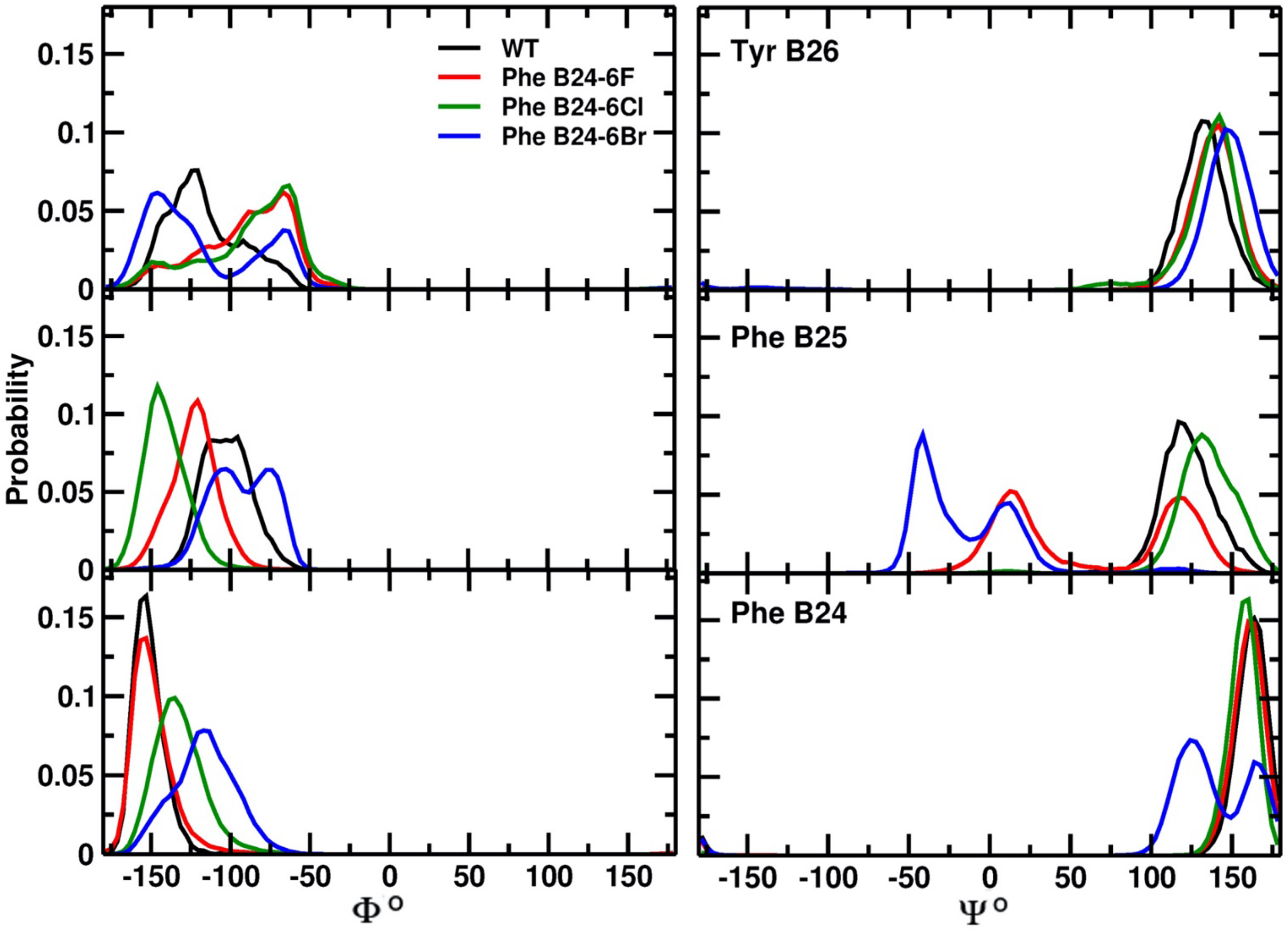


**Figure S7.** Dihedral angles distribution of Phe$^{B24}$, Phe$^{B25}$ and Tyr$^{B26}$ as a function of the halogen introduced in *ortho*-6 to Phe$^{B24}$.

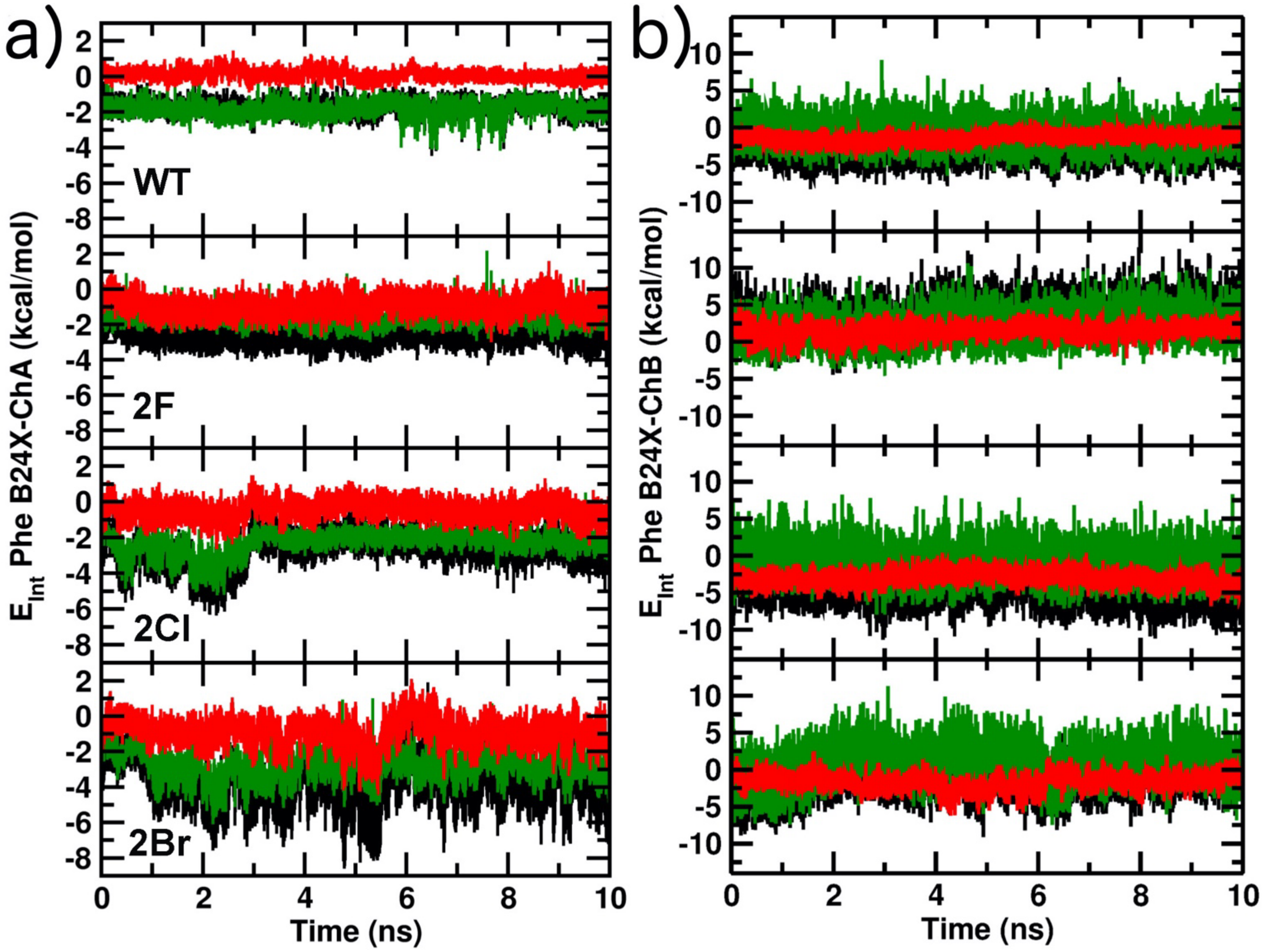


**Figure S8.** Interaction energy between: a) Phe[B24] side-chain atoms and chain A b) Phe[B24] side-chain atoms and chain B of the WT monomer, and the 2X-Phe[B24] mutants, from 10 ns of MD simulation. The total nonbonded energy is in black, the electrostatic energy contribution is in red, and the vdW energy contribution is in green.

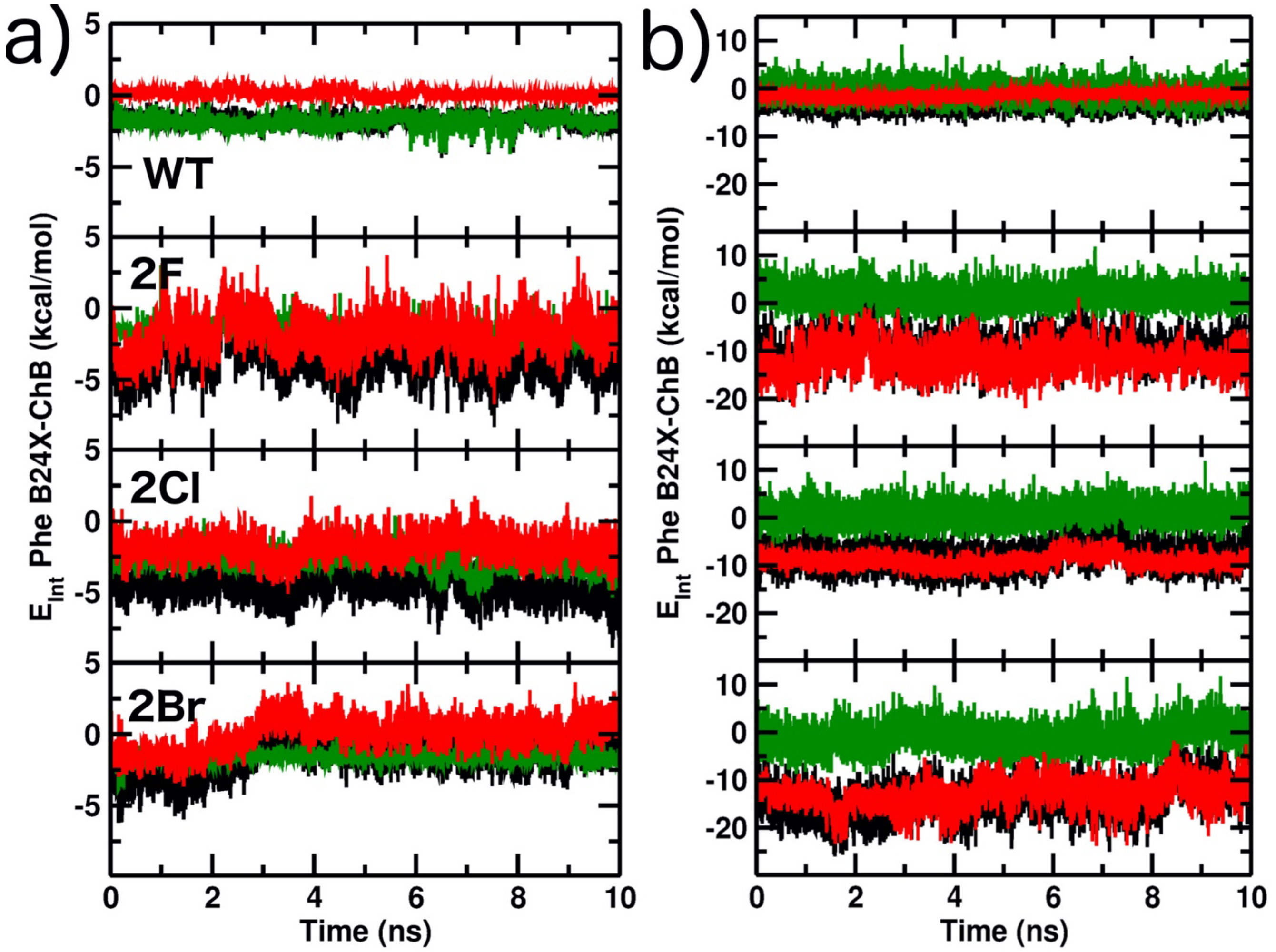


**Figure S9.** Interaction energy between: a) Phe$^{B24}$ side-chain atoms and chain A b) Phe$^{B24}$ side-chain atoms and chain B of the WT monomer, and the 6X-Phe$^{B24}$ mutants, from 10 ns of MD simulation. The total nonbonded energy is in black, the electrostatic energy contribution is in red, and the vdW energy contribution is in green.

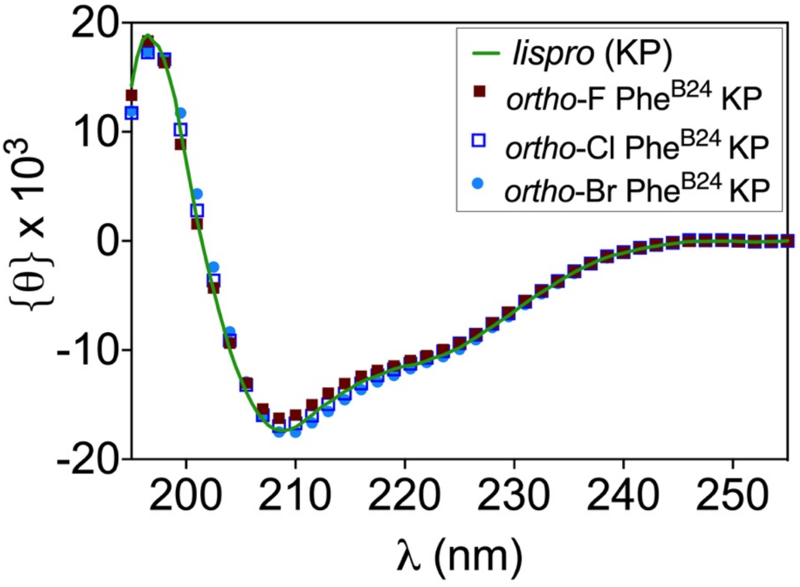


**Figure S10.** CD spectra of insulin analogs. The spectrum of the parent analog, insulin *lispro* (KP-insulin) is shown as a green line. Analogs: (■) *ortho*-F-Phe$^{B24}$-KP-insulin, (□), *ortho*-Cl-Phe$^{B24}$-KP-insulin and (●) *ortho*-Br-Phe$^{B24}$-KP-insulin. (The active ingredient of clinical product Humalog® (Eli Lilly), insulin *lispro* contains paired substitutions Pro$^{B28}$ →Lys and Lys$^{B29}$→Pro.) Spectra were sampled every 0.5 nm with 1 mm pathlength; 60% of data points are shown. The temperature was 25 °C.

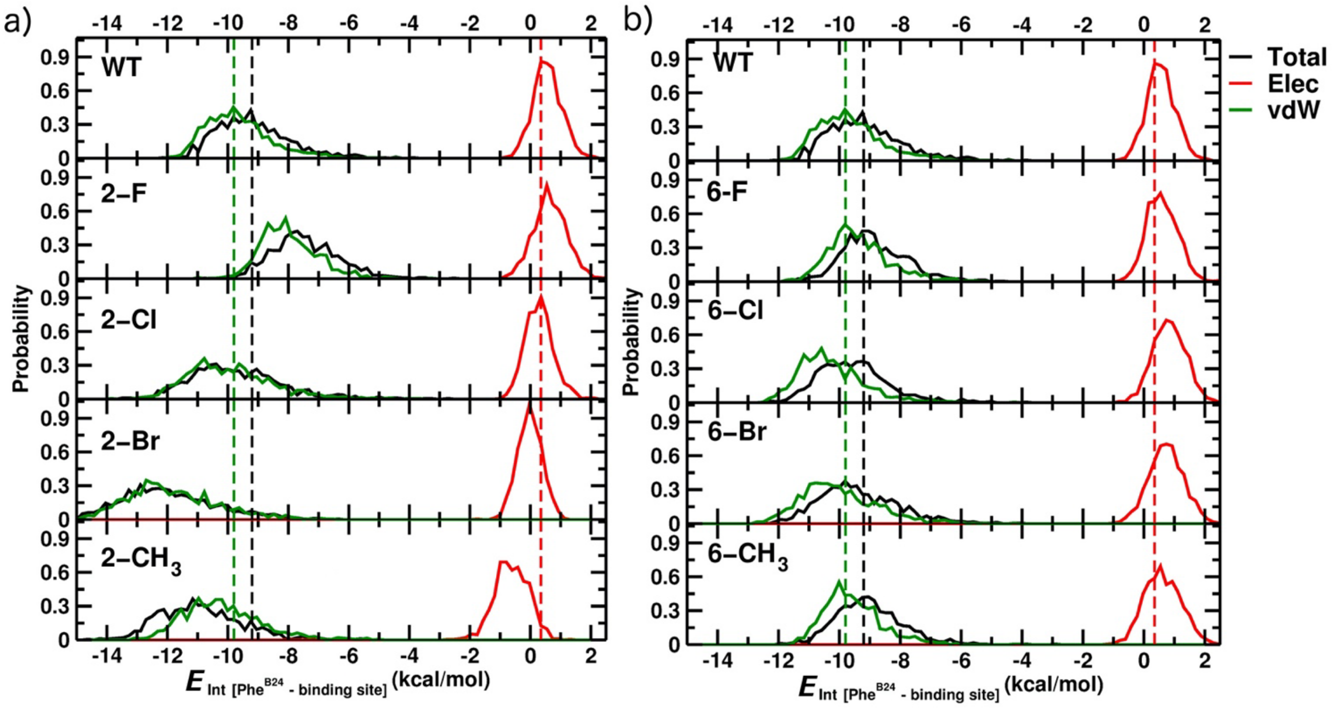


**Figure S11.** Probability distributions for the interaction energy between both units (INS monomer and micro-receptor (μIR)) from 1 ns of MD simulations for a) the 2-X and b) the 6-X halo-aromatic substitutions; $CH_3$ values are also reported as substituents of similar size but without significant electronegativity or inductive effects. Dashed lines report averaged energy values of the WT-INS bound to the μIR fragments, as averaged over the 1 ns MD simulation.

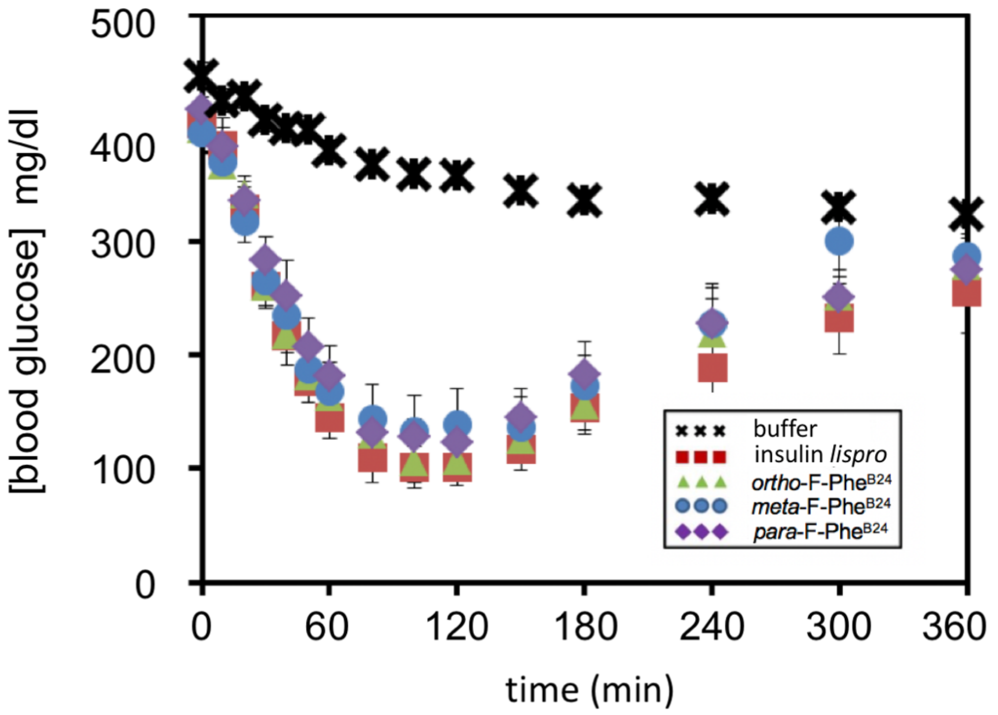


**Figure S12.** Rat studies of fluorine substituted insulin analogues. Time course of [blood glucose] following Subcutaneous (SQ) injection. Insulin lispro (■); *ortho*-F-Phe$^{B24}$-KP-insulin (▲); *meta*-F-Phe$^{B24}$-KP-insulin (●); *para*-F-Phe$^{B24}$-KP-insulin (◆); diluent buffer (╳). The doses of all insulin analogs were 2.6 nmol per 300-gram rat (n = 5).

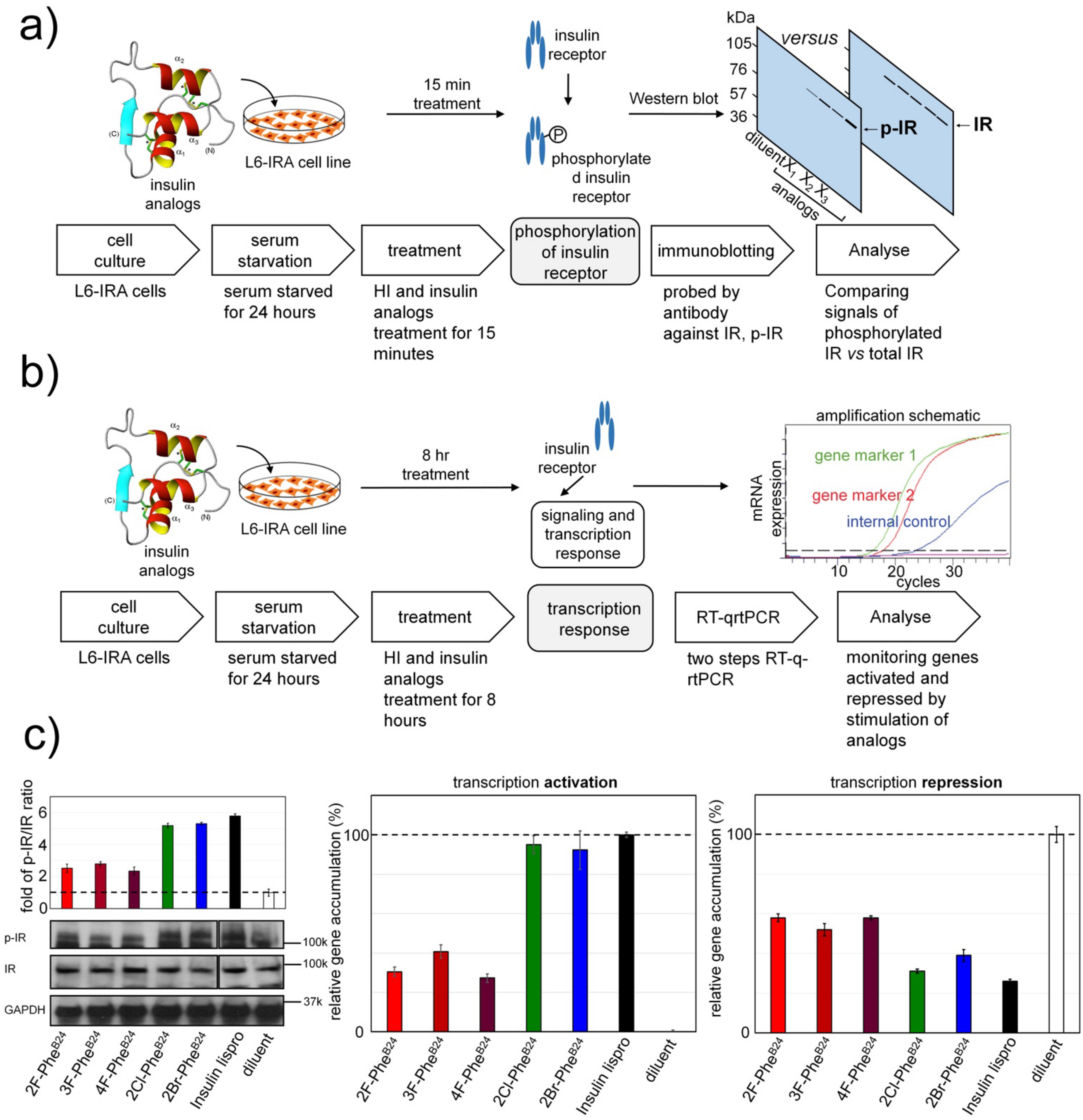


**Figure S13.** Cell-based assay of biological activities. a, b) Schematic outline of cell-based assays for the assessment of hormone-induced IR-A signaling and the transcription responses. L6 rat myoblasts stably expressing human IR isoform A (IR-A) (developed by P. De Meyts[21]) were treated with halogen analogs or control insulin lispro. After 15 min treatment Tyr-phosphorylation of IR was assessed by Western blotting [24]: (panel a). To monitor the induced transcriptional responses, L6-IRA cells treated for 8 h were collected and the transcriptional activation of cyclin D1 and repression of cyclin G2 (cell cycle schematic adapted from ref 25) assessed by rt-qPCR (schematic adapted from: Bio-Rad Labs. (2017) qPCR assay design and optimization. http://www.bio-rad.com/en-us/applications-technologies/qpcr-assay-design-optimization). The increased accumulation of cyclin D1 and decreased cyclin G2 mRNA served as readouts for insulin-induced activities (panel b). c) Histogram representation of the average fold increase over diluent from L6-IRA Western blots of p-IR/IR (left). The qPCR assay shows the increased accumulation of cyclin D1 (middle panel) as the probe of insulin-stimulated transcription activation and the decreased cyclin G2 mRNA (right panel) as the probe of insulin-stimulated transcription repression.

**Table S1.** Energy contribution differences, due to the Phe$^{B24}$-2X substitutions, from the backbone "bb" (left panels) and the side chains "sc" (right panels) atoms of the affected residues averaged over the 10 ns MD simulations.

| **ΔE $_{inter}$ (kcal/mol)** | **2F** | | | | | **2Cl** | | | | | **2Br** | | | | |
|---|---|---|---|---|---|---|---|---|---|---|---|---|---|---|---|
| | **bb contribution** | | **sc contribution** | | **Total** | **bb contribution** | | **sc contribution** | | **Total** | **bb contribution** | | **sc contribution** | | **Total** |
| | **vdW** | **elec** | **vdW** | **elec** | | **vdW** | **elec** | **vdW** | **elec** | | **vdW** | **elec** | **vdW** | **elec** | |
| **Gly A1** | -0.01 | -0.30 | 0.00 | 0.00 | **-0.31** | 0.00 | -0.69 | 0.00 | 0.00 | **-0.69** | 0.00 | -0.59 | 0.00 | 0.00 | **-0.59** |
| **Glu A17** | 0.00 | 0.05 | 0.00 | -0.33 | **-0.27** | 0.01 | 0.05 | 0.00 | -0.19 | **-0.13** | 0.00 | 0.02 | 0.00 | -0.08 | **-0.06** |
| **Val B12** | -0.08 | -0.06 | 0.03 | 0.03 | **-0.08** | 0.11 | -0.26 | 0.15 | -0.04 | **-0.03** | 0.13 | -0.15 | 0.31 | 0.00 | **0.30** |
| **Glu B13** | 0.00 | -0.15 | 0.00 | -0.32 | **-0.47** | 0.02 | -0.14 | 0.00 | -0.34 | **-0.47** | 0.03 | -0.12 | 0.01 | -0.78 | **-0.86** |
| **Leu B15** | -0.53 | -0.61 | -0.20 | -0.12 | **-1.46** | -0.20 | -0.30 | 0.27 | -0.03 | **-0.26** | -0.38 | -0.29 | -0.26 | -0.03 | **-0.97** |
| **Tyr B16** | -0.19 | 0.21 | -0.42 | -0.06 | **-0.45** | -0.04 | 0.03 | 0.32 | -0.05 | **0.26** | -0.21 | -1.26 | -0.07 | -0.17 | **-1.71** |
| **Gly B20** | -0.20 | -0.46 | 0.00 | 0.00 | **-0.66** | -0.01 | -0.96 | 0.00 | 0.00 | **-0.96** | -0.15 | -0.25 | 0.00 | 0.00 | **-0.41** |
| **Glu B21** | -0.02 | 0.00 | 0.00 | -0.83 | **-0.86** | 0.03 | -0.01 | 0.01 | -0.90 | **-0.87** | 0.01 | -0.06 | 0.00 | -0.91 | **-0.96** |
| **Gly B23** | 0.14 | 0.75 | 0.00 | 0.00 | **0.88** | 0.11 | -1.02 | 0.00 | 0.00 | **-0.91** | 0.26 | 0.58 | 0.00 | 0.00 | **0.85** |
| **Phe B24** | -0.15 | 2.03 | -0.92 | -5.32 | **-4.36** | 0.07 | -2.36 | -1.07 | -1.28 | **-4.64** | -0.09 | -3.15 | -1.60 | -1.03 | **-5.87** |
| **Phe B25** | -0.22 | -0.26 | -0.04 | 0.16 | **-0.38** | -0.07 | -0.43 | 0.00 | 0.10 | **-0.40** | 0.08 | 0.99 | 0.00 | 0.05 | **1.13** |
| **Tyr B26** | 0.02 | 0.25 | -0.52 | -0.92 | **-1.17** | 0.01 | 0.08 | 0.01 | 0.07 | **0.17** | -0.04 | -0.12 | 0.58 | -0.08 | **0.34** |
| **Total** | **-1.25** | **1.45** | **-2.08** | **-7.70** | **-9.58** | **0.05** | **-6.01** | **-0.29** | **-2.68** | **-8.94** | **-0.37** | **-4.38** | **-1.04** | **-3.02** | **-8.81** |

**Table S2.** Energy contribution differences, due to the Phe$^{B24}$-6X substitutions, from the backbone "bb" (left panels) and the side chains "sc" (right panels) atoms of the affected residues averaged over the 10 ns MD simulations.

| ΔE $_{inter}$ (kcal/mol) | 6F | | | | | 6Cl | | | | | 6Br | | | | |
|---|---|---|---|---|---|---|---|---|---|---|---|---|---|---|---|
| | bb contribution | | sc contribution | | Total | bb contribution | | sc contribution | | Total | bb contribution | | sc contribution | | Total |
| | vdW | elec | vdW | elec | | vdW | elec | vdW | elec | | vdW | elec | vdW | elec | |
| **Glu A4** | 0.00 | 0.06 | 0.00 | -1.04 | **-0.98** | 0.00 | 0.07 | 0.00 | -1.28 | **-1.21** | 0.00 | 0.06 | 0.00 | -0.76 | **-0.71** |
| **Tyr A19** | 0.05 | -0.07 | 0.12 | 0.09 | **0.19** | 0.21 | 0.07 | 0.48 | 0.13 | **0.90** | 0.07 | -0.22 | 0.02 | 0.01 | **-0.12** |
| **Cys A20** | 0.05 | -1.06 | 0.03 | 0.13 | **-0.84** | 0.13 | -0.76 | 0.05 | 0.08 | **-0.50** | 0.00 | -0.46 | 0.00 | 0.11 | **-0.36** |
| **Gln A21** | 0.01 | 1.15 | -0.05 | -0.49 | **0.63** | 0.01 | 0.84 | -0.07 | -0.27 | **0.51** | 0.00 | -0.32 | -0.02 | -0.23 | **-0.58** |
| **Phe B1** | 0.00 | -0.76 | 0.00 | 0.00 | **-0.76** | 0.00 | -0.10 | 0.00 | 0.00 | **-0.10** | 0.00 | -0.53 | 0.00 | 0.00 | **-0.53** |
| **Leu B15** | -0.11 | 1.43 | -0.03 | 0.18 | **1.46** | -0.21 | 0.23 | 0.30 | 0.08 | **0.40** | -0.35 | -0.37 | -0.18 | -0.04 | **-0.94** |
| **Tyr B16** | -0.05 | 0.26 | -0.23 | -0.84 | **-0.86** | -0.17 | -0.01 | 0.02 | -0.66 | **-0.82** | -0.23 | 0.02 | -0.19 | 0.08 | **-0.32** |
| **Val B18** | -0.01 | -0.37 | 0.00 | 0.06 | **-0.32** | -0.02 | -0.20 | 0.01 | 0.03 | **-0.18** | 0.01 | -0.33 | 0.00 | 0.01 | **-0.31** |
| **Cys B19** | -0.20 | -0.93 | -0.12 | -0.48 | **-1.72** | -0.25 | -0.50 | -0.14 | -0.09 | **-0.98** | -0.14 | -0.70 | 0.22 | -0.07 | **-0.68** |
| **Arg B22** | -0.03 | -0.46 | 0.01 | -2.18 | **-2.66** | -0.02 | -0.18 | 0.01 | -1.39 | **-1.58** | 0.05 | -0.37 | 0.04 | -0.77 | **-1.05** |
| **Gly B23** | 0.05 | -1.65 | 0.00 | 0.00 | **-1.60** | 0.06 | -1.65 | 0.00 | 0.00 | **-1.59** | 0.37 | -0.90 | 0.00 | 0.00 | **-0.53** |
| **Phe B24** | 0.22 | 4.11 | -1.02 | 1.98 | **5.29** | 0.27 | 4.17 | -1.25 | 2.24 | **5.42** | 0.53 | 4.46 | -2.10 | 2.64 | **5.53** |
| **Phe B25** | -0.24 | -1.61 | -0.03 | -0.23 | **-2.11** | 0.10 | -1.37 | 0.08 | -0.16 | **-1.35** | -0.17 | -0.46 | -0.02 | -0.20 | **-0.87** |
| **Tyr B26** | -0.09 | -1.35 | -0.44 | 0.52 | **-1.36** | 0.03 | -1.03 | -0.42 | 0.21 | **-1.22** | 0.18 | -0.95 | -0.22 | 0.49 | **-0.50** |
| **Thr B30** | 0.00 | -1.44 | 0.00 | 0.00 | **-1.44** | 0.00 | -0.86 | 0.00 | 0.00 | **-0.85** | 0.01 | -0.73 | 0.00 | -0.01 | **-0.72** |
| **Total** | **-0.36** | **-2.68** | **-1.76** | **-2.28** | **-7.08** | **0.02** | **4.94** | **-0.86** | **3.90** | **-3.15** | **0.32** | **-1.79** | **-2.45** | **1.25** | **-2.67** |

**Table S3**. Properties of insulin analogs.

| *analog* | $\Delta G_u$[a] (kcal/mol) | $\Delta\Delta G_u$ (kcal/mol) | *m*-value[b] [M GuHCl] | $C_{mid}$[c] (kcal/mol/M) | Receptor binding hIR-B (nM) ± SD |
|---|---|---|---|---|---|
| KP-insulin | 2.8 ± 0.1 | - | 0.61 ± 0.01 | 4.6 ± 0.1 | 0.09 ± 0.01 (89%) |
| *ortho*-F-Phe$^{B24}$ | 3.8 ± 0.1 | -1.0 ± 0.1 | 0.76 ± 0.01 | 5.0 ± 0.1 | 0.47 ± 0.07 (17%) |
| *meta*-F-Phe$^{B24}$ | 3.0 ± 0.1 | -0.2 ± 0.1 | 0.67 ± 0.01 | 4.5 ± 0.1 | 0.26 ± 0.04 (30%) |
| *para*-F-Phe$^{B24}$ | 2.75 ± 0.10 | 0.05 ± 0.1 | 0.62 ± 0.01 | 4.5 ± 0.1 | 0.13 ± 0.02 (62%) |
| *ortho*-Cl-Phe$^{B24}$ | 3.8 ± 0.1 | -1.0 ± 0.1 | 0.77 ± 0.01 | 4.9 ± 0.1 | 0.22 ± 0.03 (36%) |
| *ortho*-Br-Phe$^{B24}$ | 3.6 ± 0.1 | -0.8 ± 0.1 | 0.75 ± 0.01 | 4.8 ± 0.1 | 0.26 ± 0.04 (31%) |

[a] Data are from two-state modeling of CD-guanidine titrations performed at 25 °C and pH 7.4.
[b] The *m*-value (units of kcal/(mol/M)) is the slope of unfolding free energy $\Delta G_u$ versus molar concentration of denaturant.
[c] $C_{mid}$ is the guanidine denaturant concentration at which 50% of the protein is in the unfolded state.

**Table S4.** Fibrillation data for the halogenated insulin analogs. *p*-values are reported in Supplemental Table S5.

| Analog | Accelerated protocol hr ± SD (n) [a] | Gentle rocking protocol Days ± SD (n) [a] |
|---|---|---|
| KP-ins | 9.7 ± 2.6 (8) | 4.1 ± 1.0 (3) |
| 2F-Phe$^{B24}$-KP-ins | 16.6 ± 1.8 (8) | 9.7 ± 2.9 (3) |
| 3F-Phe$^{B24}$-KP-ins | 8.3 ± 0.6 (8) | 12.3 ± 0.6 (3) |
| 4F-Phe$^{B24}$-KP-ins | 7.1 ± 1.1 (8) | 12.0 ± 1.0 (3) |
| 2Cl-Phe$^{B24}$-KP-ins | 15.5 ± 1.2 (8) | 15.7 ± 2.5 (3) |
| 2Br-Phe$^{B24}$-KP-ins | 21.6 ± 1.9 (8) | 12.0 ± 1.0 (3) |

[a] The number of replicates is given by (n).

**Table S5**. *p*-values for the fibrillation data of halogenated insulin analogs.

| analog | *p*-value[a] | |
|---|---|---|
| | Accelerated protocol | Gentle rocking protocol |
| KP-ins | - | - |
| 2F-Phe$^{B24}$-KP-ins | 0.00001 | 0.01706 |
| 3F-Phe$^{B24}$-KP-ins | 0.07826 | 0.00013 |
| 4F-Phe$^{B24}$-KP-ins | 0.00929 | 0.00031 |
| 2Cl-Phe$^{B24}$-KP-ins | 0.00002 | 0.00086 |
| 2Br-Phe$^{B24}$-KP-ins | 2.56E-08 | 0.00031 |

[a] *p*-value <0.05 is considered as significantly different.
